# Targeting G9a translational mechanism of SARS-CoV-2 pathogenesis for multifaceted therapeutics of COVID-19 and its sequalae

**DOI:** 10.1101/2024.03.04.583415

**Authors:** Adil Muneer, Ling Xie, Xuping Xie, Feng Zhang, John A. Wrobel, Yan Xiong, Xufen Yu, Charles Wang, Ciprian Gheorghe, Ping Wu, Juan Song, Guo-Li Ming, Jian Jin, Hongjun Song, Pei-Yong Shi, Xian Chen

**Author notes:** These authors contributed equally to this work (Adil Muneer, Ling Xie).

## Abstract

By largely unknown mechanism(s), SARS-CoV-2 hijacks the host translation apparatus to promote COVID-19 pathogenesis. We report that the histone methyltransferase G9a noncanonically regulates viral hijacking of the translation machinery to bring about COVID-19 symptoms of hyperinflammation, lymphopenia, and blood coagulation. Chemoproteomic analysis of COVID-19 patient peripheral mononuclear blood cells (PBMC) identified enhanced interactions between SARS-CoV-2-upregulated G9a and distinct translation regulators, particularly the N^6^-methyladenosine (m^6^A) RNA methylase METTL3. These interactions with translation regulators implicated G9a in translational regulation of COVID-19. Inhibition of G9a activity suppressed SARS-CoV-2 replication in human alveolar epithelial cells. Accordingly, multi-omics analysis of the same alveolar cells identified SARS-CoV-2-induced changes at the transcriptional, m^6^A-epitranscriptional, translational, and post-translational (phosphorylation or secretion) levels that were reversed by inhibitor treatment. As suggested by the aforesaid chemoproteomic analysis, these multi-omics-correlated changes revealed a G9a-regulated translational mechanism of COVID-19 pathogenesis in which G9a directs translation of viral and host proteins associated with SARS-CoV-2 replication and with dysregulation of host response. Comparison of proteomic analyses of G9a inhibitor-treated, SARS-CoV-2 infected cells, or *ex vivo* culture of patient PBMCs, with COVID-19 patient data revealed that G9a inhibition reversed the patient proteomic landscape that correlated with COVID-19 pathology/symptoms. These data also indicated that the G9a-regulated, inhibitor-reversed, translational mechanism outperformed G9a-transcriptional suppression to ultimately determine COVID-19 pathogenesis and to define the inhibitor action, from which biomarkers of serve symptom vulnerability were mechanistically derived. This cell line-to-patient conservation of G9a-translated, COVID-19 proteome suggests that G9a inhibitors can be used to treat patients with COVID-19, particularly patients with long-lasting COVID-19 sequelae.

## INTRODUCTION

The Coronavirus Disease (COVID-19) pandemic, caused by severe acute respiratory syndrome coronavirus-2 (SARS-CoV-2), is an unprecedented global public health crisis having already claimed more than 6.8 million lives worldwide.^1^ Particularly, high mortality is observed for SARS-CoV-2-infected patients who have pre-existing chronic conditions such as recovery from sepsis, chronic pulmonary diseases, metabolic diseases (e.g., diabetes), asthma, cardiovascular diseases, thrombosis, chronic liver disease and cirrhosis, and cancer.^2, 3^ Despite initial success of vaccines in reducing COVID-19 infections, hospitalizations, and deaths, neutralizing antibody levels eventually wane with time ^4, 5^ and genomic variation owing to low replication fidelity^6, 7^ has led to emergence of SARS-CoV-2 variants with increased transmissibility or virulence. For example, an Omicron subvariant, XBB.1.16, also known as Arcturus, has fueled a surge of COVID-19 cases in 2022^8^. Accordingly, none of the available monoclonal antibodies, with emergency use authorization, neutralize omicron and its variants effectively^9–11^. Also, development of drug resistance has been observed for antiviral drugs that target either SARS-CoV-2 polymerase (e.g., Remdesivir, Molnupiravir) or protease (e.g. Paxlovid)^12–14^. More importantly, without available clinical drugs, emerging cases of post-acute sequela of COVID-19 (aka “long COVID”) have been reported^15^. For example, COVID-19 causes a plethora of neurological, neuropsychiatric, and psychological impairments such as ischemic and hemorrhagic stroke, encephalopathy, encephalitis, brain fog, depression, anxiety, and sleep disorders^16^. Yet, SARS-CoV-2 pathogenesis and the etiology of long COVID neurological symptoms are poorly understood. To effectively combat both emerging variants of the coronavirus and long-lasting COVID-19 sequelae, key mechanistic questions need to be answered, including (1) how the SARS-CoV2 replication/life cycle is regulated in the host, and (2) how SARS-CoV2 hijacks host pathways to promote COVID-19 pathogenesis.

The molecular/cellular hallmarks of COVID-19 pathogenesis include increased proportion of monocyte-derived macrophages, reduction and functional exhaustion of T-cells (lymphopenia), and increased levels of serum cytokines (hyperinflammation);^17, 18^ together, these circumstances result in sepsis and acute respiratory distress syndrome (ARDS), two leading complications associated with severe COVID-19^19, 20^. In addition, in correlation with the crucial function of histone modification and chromatin remodeling during SARS-CoV-2 infection^21^, the histone methyltransferases G9a and G9a-like protein (GLP; hereafter G9a will represent both proteins) showed upregulated expression in COVID-19 patients who had high virus load,^22^ implicating G9a-associated pathways and mechanisms in COVID-19 pathogenesis. However, the well-understood canonical function of G9a in gene-specific transcriptional silencing did not explain how translation of other proteins was upregulated in the immunocompromised states of COVID-19.

Proteomic dissection of endogenous protein-protein interaction complexes (interactomes) has unique strengths for discovery of new functions of bait proteins, which can be extrapolated by identifying interactors that have known functions^23, 24^. Because the systemic cytokine profiles in severe COVID-19 patients were similar to profiles in macrophage activation syndromes^25^, particularly viral sepsis^26^, we used our chromatin activity-based chemoproteomic (ChaC) with a biotinylated G9a inhibitor UNC0965^27^ to dissect G9a-interacting pathways in the peripheral blood mononuclear cells (PBMCs) of COVID-19 patients. Notably, unlike conventional immunoprecipitation (IP)-MS^23, 24^ that characterizes protein complexes based only on epitope abundance, ChaC identified *in vivo* interactors of constitutively active G9a specifically from the diseased cells in the PBMCs with different cell types. Akin to what was found in endotoxin-tolerant (ET) macrophages ^28^ that have molecular characteristics similar to the immunopathological background of COVID-19 vulnerable groups that have pre-existing chronic inflammatory diseases^17, 18^, UNC0965 ChaC-MS identified numerous regulators of translation, ribosome biogenesis, and proteostasis that had enhanced interaction with G9a in COVID-19 patient PBMCs. Coincidently, in the lungs of deceased COVID-19 patients, major protein translation pathways were found dysregulated^29^. In addition, in a recent study of virus-host protein interactome, Zhou et al. revealed that, via interactions with host translational regulators, SARS-CoV-2 hijacks the corresponding translational pathways^30^. Thus, our ChaC identification of G9a interactions with multiple regulators of SARS-CoV-2 hijacked translation now extend G9a function beyond its transcriptional repression function to translational regulation of SARS-CoV-2 pathogenesis.

Specifically, in COVID-19 related samples, G9a-interacting translation regulators included the N^6^-methyladenosine (m^6^A) RNA methylase METTL3,^31, 32^ the ribosomal (r)RNA methylase fibrillarin (FBL), and another histone methyltransferase Ezh2. METTL3 is implicated in viral m^6^A RNA modification and SARS-CoV2 dysregulated host immune response^33–36^, and elevated m^6^A levels were found associated with severe clinical outcomes and mortality of COVID19 patients^37–40^. SARS-CoV-2 can escape the host cell innate immune response by mimicking the host mRNA capping machinery and 2′-O methylation (2′-O-Me), the most prevalent modification in rRNA; FBL is the only known methyltransferase that catalyzes site-specific 2′-O-Me of rRNA. Importantly, Ezh2 and FBL, both of which interact with G9a, were found in a complex that regulates the 2′-O-Me rRNA-mediated protein synthesis^41^. The abovementioned reports in conjunction with our ChaC findings raised a triggering possibility that, via COVID19-characteristic interactions with METTL3 or Ezh2/FBL, G9a coordinates m^6^A– or Ezh2-mediated SARS-CoV-2 pathogenesis.

Bojkova et al. reported that viral replication was prevented by inhibitors of translational pathways reshaped by SARS-CoV-2^42^. We showed that G9a and Ezh2 inhibitors suppressed viral replication in SARS-CoV-2 infected human alveolar epithelial cells that overexpressed human ACE2 (A549-hACE2). Accordingly, we used G9a and Ezh2 inhibitors as the mechanistic probes to conduct COVID-19 pathology-correlated, multiomics analyses to investigate inhibitor effects on the SARS-CoV-2 infection-induced transcriptome, m^6^A RNA epitranscriptome, proteome, phosphoproteome, and secretome of the A549-hACE2 cells or *ex vivo* culture of COVID-19 patient PBMCs. Correlated analyses of these multiomics data showed that G9a inhibition reversed SARS-CoV-2-induced changes in m^6^A RNA abundance and/or expression, phosphorylation, or secretion of specific proteins; these proteins whose translation was regulated by G9a (i.e., affected by G9a inhibitor) unite the networks associated with major stages of COVID-19 pathogenesis including host-virus interactions, and dysregulated host immune response. These results indicated that constitutively active G9a is the upstream/master regulator of widespread translational or post-translational (e.g., phosphorylation or secretion) processes associated with COVID-19 pathogenesis. Further, these results elucidated the mechanisms of inhibitor action toward both virus and the dysregulated host response that broadly reversed synthesis and degradation of specific proteins that ultimately define COVID-19 pathology. In correlation with the fact that abnormal m^6^A modification enhances the replication of a broad range of coronaviruses/variants and their associated pathogenesis^43, 44^, we found that G9a inhibition reversed the m^6^A epitranscriptome landscape associated with SARS-CoV2 infection. These results implicated G9a inhibition in broad-spectrum blockade of emerging coronavirus variants or any virus. In addition, we identified multiple G9a-regulated, inhibitor-reversed pathways characteristic of long COVID or COVID neurological symptoms. Importantly, akin to our finding that a brain-penetrant inhibitor of G9a, MS1262, reversed brain neuropathological pathways associated with Alzheimer’s disease pathogenesis^45^, we found that MS1262 suppressed SARS-CoV2 replication, suggesting a therapeutic effect of G9a inhibition on long-lasting COVID neurological symptoms.

In sum, our COVID-19 pathology correlated multi-omics studies reveal a novel G9a-translation mechanism of COVID-19 pathogenesis from which we derived/validated biomarkers that can identify patient populations vulnerable to severe symptoms. Accordingly, targeting G9a and its interactor Ezh2 represents both virus– and host-directed therapeutics of severe COVID-19 and long COVID.

## RESULTS

### G9a interactome is implicated in the translational regulation of COVID-19 pathogenesis

To determine at which regulatory layer G9a promotes COVID-19 pathogenesis, we dissected G9a-associated pathways by our label-free quantitation (LFQ)^46^ ChaC-MS approach. We used two ChaC probes, i.e., UNC0965 for G9a^27^ and UNC2399 for Ezh2^47^ for capturing G9a– and Ezh2-interacting complexes in the peripheral blood mononuclear cells (PBMCs) from COVID-19 patients (RayBiotech Life). For comparison, we included ChaC-MS results for G9a complexes obtained from ET macrophage cells^28^ (**Figure 1a**). Three technical replicates were performed for each biological replicate, and principal component analysis (PCA) showed good separation between G9a, EZH2, and control (UNC0125) pulldowns from patient PBMCs (**Figure S1c**). Based on LFQ ratios that are proportional to the relative binding of individual proteins to G9a or Ezh2 in COVID-19 patient PBMCs, LFQ ChaC-MS identified 1319 protein groups, of which 822 and 410 proteins were identified as G9a and Ezh2 interactors, respectively, in at least one COVID19 patient, and 368 proteins shared between the two methyltransferases. (**Figure 1b & S1d; Supplemental Table S1a**). Unsurprisingly, there was significant overlap between G9a/Ezh2 interactors in COVID19 patients and ET macrophages. In agreement with the canonical epigenetic (transcriptional) regulatory function of G9a and EZH2, we identified several proteins associated with chromatin remodeling and histone modification, including the SWI/SNF remodeling complex and BRD4, which were identified by siRNA and genome-wide CRISPR screens as essential host factors for SARS and pan-coronavirus infection, respectively ^21, 48^ (**Figure 1b & S1e**). Similarly, in line with reports that core components of the G9a complex including EHMT1 and WIZ interact with SARS-CoV-2 encoded ORF9c ^49, 50^, we found 51 ChaC-identified interactors associated with ORF9c, and turnover of 24 other proteins was upregulated following infection^42^ (**Figure 1b; Supplemental Table S1b**), indicating the regulatory function of G9a interactome in COVID-19 immunopathogenesis.

**Figure 1.**
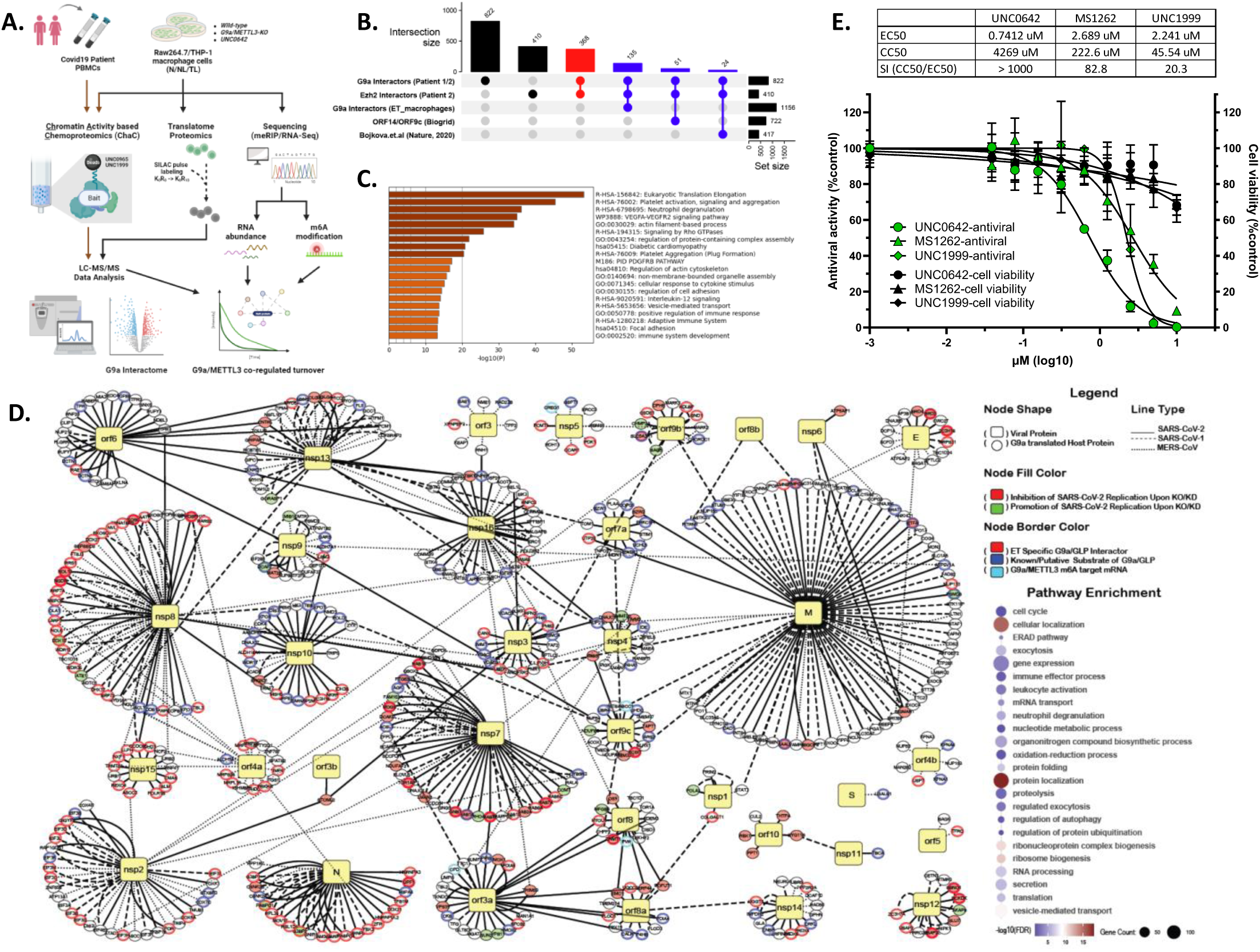
G9a interacts with host translation regulators to promote SARS-CoV-2 replication and proviral gene expression. (**A**) Schematic overview of multi-omics workflow to dissect function of G9a during COVID-19. **(B)** Upset plot showing number of ChaC-identified G9a/EZH2 interactors in patient PBMCs or ET macrophage cells. Shared interactors are shown in red; in blue is shown interactor overlap with ChaC-MS results from ET macrophages, ORF14 host interactors and proteins with SARS-CoV-2 upregulated turnover. The histogram on right shows the total number of hits in each dataset. **(C)** Pathway enrichment results for 368 G9a/EZH2 shared interactors identified from patient PBMCs. **(D)** Virus-host protein-protein interaction map depicting G9a-translated host interactors. Human proteins are shown as circles, whereas viral proteins are represented by yellow squares. Each edge represents an interaction between a human and a SARS-CoV-2 (solid line), SARS-CoV-1 (dashed line) or MERS-CoV (dotted line) protein with certain interactions shared between these three viruses. Node border colors show ET/PBMC-specific G9a interactors that are, respectively, G9a-translated host proteins (red), G9a/GLP substrates (blue) or G9a/METTL3-coregulated m^6^A mRNAs (cyan). Node fill color shows that genetic perturbation of several of these G9a-translated proteins hinders (red) or promotes (green) SARS-CoV-2 replication/infection. Pathway enrichment scores for 503 G9a translated host interactors are shown on the side. ET-specific G9a interactors and G9a/METTL3-coregulated m^6^A targets were defined in [25], whereas host-virus physical interactions, effects of genetic perturbation on SARS-CoV-2 replication/infection, and G9a/GLP substrate definitions were curated manually from literature sources. **(E)** Antiviral activity (n = 2) and cytotoxicity (n = 3) of indicated compounds were evaluated in A549-hACE2 cells. Half-maximum effective concentration (EC_50_) and cytotoxicity concentration (CC_50_) values were calculated by fitting a nonlinear regression model (four parameters) with results summarized. (See also **Figures S1 & S2, Supplemental Table S1**).

In addition, ChaC revealed that G9a and Ezh2 interacted with the same translational regulators in ET macrophages and COVID-19 patient PBMCs (**Figure 1b-1c & S1e**). Notably, among the ChaC-identified G9a-interacting translation regulators, the splicing factor SF3B1 and the 40S ribosomal protein Rps14 were identified as patient PBMC– and ET-specific G9a interactors (**Supplemental Table S1b**); Bojkova et al. showed that emetine inhibition of Rps14 and pladienolide inhibition of SF3B1 significantly reduced SARS-CoV-2 replication^42^. In correlation with the newly characterized function of EZH2 in translation regulation^41^, numerous members of the EZH2 complex were identified, such as EZH1, EZH2, EED, and SUZ12. Broadly, interactors shared between patient PBMC– or ET-macrophage specific G9a and EZH2 were primarily involved in translation elongation, immunity, angiogenesis and blood coagulation, cytoskeletal rearrangement and vesicle-mediated transport – all pathways that are dysregulated by SARS-CoV-2 infection ^19, 51–56^ (**Figure 1c**). These UNC0965 ChaC findings were highly correlated with COVID-19 clinicopathology because 1) in nasopharyngeal swabs and/or autopsy samples from severe COVID-19 patients, multiple G9a, EZH2, and METTL3 complex members are observed to be simultaneously dysregulated (**Figure S1a-S1b**), 2) histone mimicry by SARS-CoV-2 encoded ORF8 resulted in simultaneous increases of G9a– and Ezh2-catalyzed methylation at the histone H3 K9 and K27 following SARS-CoV-2 infection^57, 58^, and 3) EZH2 binds METTL3^59^ and EZH2 regulates IRES-dependent translation^60^. Taken together, these SARS-CoV-2 pathology-correlated findings of ChaC-MS suggested that, via interactions with host translation regulators such as METTL3 and EZH2, SARS-CoV-2-upregulated G9a noncanonically function in the SARS-CoV-2 hijacked translational regulation of COVID immunopathogenesis.

### G9a-interacting translation pathways promote SARS-CoV-2 infection and replication

In ET macrophages that mimic the immunopathological phenotype of COVID-19, we found^28^ that G9a and METTL3 coregulated the m^6^A-mediated translation of proteins associated with sepsis and ARDS complications of severe COVID-19.^19, 20^ Specifically, our translatome proteomics and m^6^A RIP-Seq analysis of ET macrophages identified proteins whose translation was G9a-dependent (‘G9a-translated’ proteins); these proteins included 503 host interactors of SARS-CoV-1/2– and MERS-CoV-encoded proteins,^61, 62^ 11 known COVID-19 markers,^51, 52^ and 66 other coronavirus pathogenesis-related proteins (**Figure 1d & S2a-S2c;** also see **Supplemental Table S1c)**. Certain G9a-translated proteins were ChaC-identified G9a interactors and/or non-histone G9a substrates and/or G9a/METTL3-coregulated m^6^A targets in both patient PBMCs and ET macrophages (**Figure 1d**), which further supported the translational or post-translational function of G9a in COVID-19 pathogenesis. More importantly, genetic perturbation of numerous G9a-translated host interactors of SARS-CoV-1/2– and MERS-CoV-encoded proteins impairs SARS-CoV-2 replication and infection^21, 62^. These G9a-translated, host-virus interactors were primarily involved in multiple signaling pathways related to gene expression, cell cycle, immunity (e.g., leukocyte activation, neutrophil degranulation), translation (e.g., ribosome/ribonucleoprotein biogenesis), RNA processing (e.g., splicing and transport), and proteostasis (e.g., proteolysis, ubiquitination, autophagy, secretion, exocytosis, protein folding/localization/transport). In fact, all of these pathways are implicated in the SARS-CoV-2 replication/life cycle and COVID-19 pathogenesis^19, 51–56^ (**Figure 1d**). Similarly, several host factors critical for SARS-CoV-2 infection identified in siRNA/CRISPR based screens are closely related to G9a complex^21, 62^ (**Figure S2d**). These results systematically revealed the associations of G9a interactome and G9a-translated proteins with host-virus interactions and SARS-CoV-2 infection/replication.

To validate the in vivo function of the G9a ChaC-dissected interactome in SARS-CoV-2 pathogenesis, we evaluated multiple G9a– and EZH2-targeting compounds for their antiviral activity against SARS-CoV-2-Nluc in a human alveolar epithelial cell line that overexpresses human ACE2 receptor (A549-hACE2). Briefly, compounds targeting G9a including UNC0642 (EC_50_ = 0.74 µM; CC_50_ = 4269 µM; SI > 1000) and MS1262 (EC_50_ = 2.69 µM; CC_50_ = 223 µM; SI = 83) showed potent antiviral activity against SARS-CoV-2 with good selectivity indices (SI = CC_50_/EC_50_). Similarly, Ezh2 inhibitor UNC1999 (EC_50_ = 2.241 µM; CC_50_ = 48.54 µM; SI =20) hindered SARS-CoV-2 replication in this model system, whereas tazemetostat (EC_50_ = >10 µM; CC_50_ = >10 µM; SI = >1), a potent EZH2 inhibitor approved for treatment of epithelioid sarcoma and follicular lymphoma ^63^, showed moderate decrease in virus proliferation (**Figure 1e & S1f).** Thus, drugs that target G9a or its interactor Ezh2 are potent suppressors of SARS-CoV-2 replication at sub/low micromolar concentrations and with good selectivity, making them attractive candidates for COVID-19 therapy.

### SARS-CoV-2 and G9a co-regulates host response pathways primarily at the translational and post-translational levels

To further clarify the ChaC-derived function of G9a in the translational regulation of SARS-CoV-2 pathogenesis, we investigated G9a inhibitor effects on the SARS-CoV-2 infection-induced transcriptome, m^6^A RNA epitranscriptome, proteome, phosphoproteome, and secretome. Correspondingly, we conducted COVID-19 pathology-correlated, multiomics analyses of SARS-CoV-2 infected A549-hACE2 cells or *ex vivo* culture of COVID-19 patient PBMCs with and without UNC0642 (a G9a inhibitor) treatment (**Figure 2a & S3a**). Principal component analysis showed clear separation between mock– and SARS-CoV-2 infected A549-hACE2 cells while, expectedly, UNC0642 treatment led to a distinct omics landscape compared with controls (**Figure S3b**). High reproducibility was observed between biological replicates and across experimental conditions across datasets (**Figure S3c**). In total, high-quality quantification for 49,021 human or viral entities, including 29,756 transcripts, 7,461 protein families, 11,217 phosphoproteins, and 587 secreted proteins, was obtained across datasets. There were 3,211 entries (1,393 RNA-Seq, 109 proteome, 1,326 phospho-proteome and 383 secretome) that showed differential enrichment for indicated comparisons **(Figure S3d-S3e; Supplemental Table S2)**. Effective infection of A549-hACE2 by SARS-CoV-2 was confirmed by a dramatic increase in viral transcripts and encoded proteins **(Figure 2b)**.

**Figure 2.**
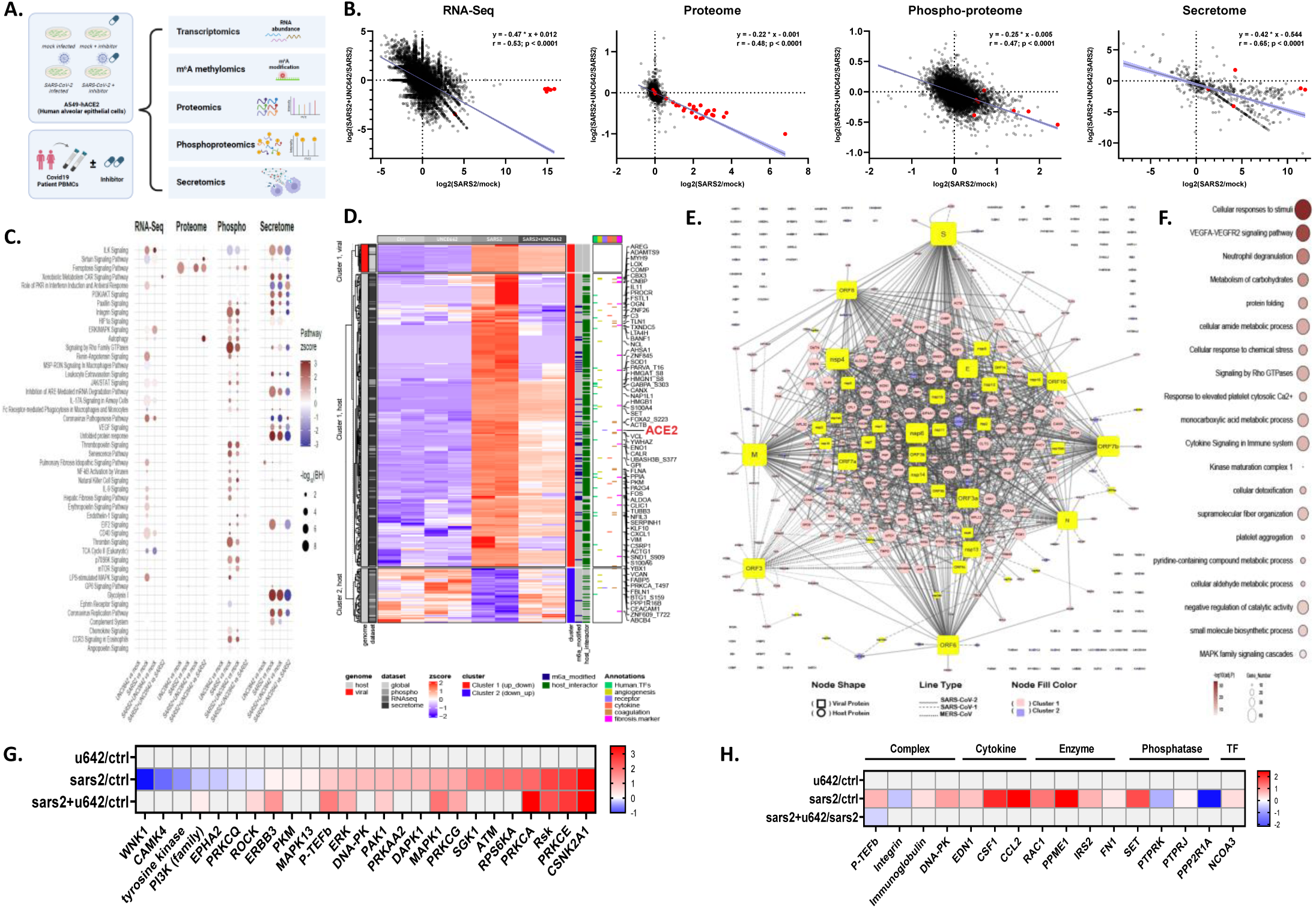
G9a inhibition reverses multi-omic landscape of SARS-CoV-2 infection. (**A**) Schematic design of multi-omic approaches to dissect G9a regulated pathogenesis of COVID-19 in A549-hACE2 cells and COVID-19 patient-derived PBMCs. **(B)** Scatter plots showing that UNC0642 treatment reverses SARS-CoV-2 induced changes to host (black) and viral (red) entities at each omics level in virus-infected A549-hACE2 cells. Linear regression (with 95% CI), slope of regression line (β), and Pearson correlation (r) are overlaid. **(C)** Plot summarizing pathway activation scores estimated using entities that are differentially regulated at indicated omics levels following SARS-CoV-2 infection and/or UNC0642 treatment of A549-hACE2 cells. Each column corresponds to indicated pairwise comparison. Red represents pathway activation, and blue represents pathway inhibition. The dot size corresponds to adjusted p-values. **(D)** Heatmap of SARS-CoV-2 dysregulated transcripts (52), proteins (23), phospho-sites (140), and secreted (237) proteins in A549-hACE2 cells whose levels are reversed by UNC0642 treatment. Viral and host entities are clustered separately and further subdivided into two groups, i.e., cluster-1 = upregulated following infection and downregulated upon UNC0642 treatment; cluster-2 = downregulated upon infection and upregulated upon UNC0642 treatment. Annotations on right highlight G9a-regulated m^6^A-modified transcripts and host interactors of SARS-CoV-1/2– and MERS-CoV-encoded proteins. Six functional clusters include viral receptors and proteases, transcription factors, cytokines/chemokines, coagulation system-related proteins, angiogenesis-associated proteins, and fibrosis markers. **(E)** Virus-host interactome map showing that nearly all cluster-1 proteins identified in (D) are host interactors of SARS-CoV-2 (solid line), SARS-CoV-1-(dashed line) or MERS-CoV-(dotted line) encoded proteins. Viral proteins are depicted by yellow rectangles, and host interactors are represented by circles. Size of each node represents connectivity, and node fill color (cluster-1 in pink; cluster-2 in blue) represents coronavirus-dysregulated/inhibitor-reversed proteins. **(F)** Gene Ontology enrichment analysis for proteins shown in (D-E). All terms with significant over-representation (adjusted P < 0.05) are kept, and redundant terms are removed. Dot size represents the number of dysregulated genes/proteins belonging to said term. **(G)** Kinase activity scores (-log10(P) < 0.05) determined based on differentially regulated phospho-sites identified in A549-hACE2 cells following SARS-CoV-2/mock infection with or without UNC0642 treatment. Rows represent indicated pairwise comparison, and columns correspond to individual kinase/family (red for activation, blue for deactivation). **(H)** Upstream regulator activity (-log10(P) ≤ 0.05; activation-score ≥ |0.5|) was estimated using differentially regulated phospho-sites identified in A549-hACE2 cells following SARS-CoV-2/mock infection with or without UNC0642 treatment. Rows represent indicated pairwise comparison, and columns correspond to individual regulator/family (red for activation, blue for deactivation). (See also **Figure S3 & S4, Supplemental Table S2**).

After cellular entry mediated by receptors and proteases, SARS-CoV-2 hijacks host translation machinery to induce a hyper-inflammatory response mediated by particular transcriptional factors (TFs). This response leads to a ‘cytokine-storm’ that is implicated in blood hypercoagulability, fibrosis and micro-thrombosis in severe COVID-19 patients^64, 65^. Correspondingly, we identified these process-related proteins that showed SARS-CoV-2-induced expression changes, including various viral receptors and proteases, TFs, cytokines (and their receptors), and proteins associated with coagulation, angiogenesis, and fibrosis (**Figure S3e**).

From a systems view, our multiomics-correlated data showed that SARS-CoV-2 infection led to activation of pathways related to (1) viral replication/life cycle – including coronavirus replication and pathogenesis, phagocytosis, xenobiotic metabolism, and LPS-stimulated MAPK signaling, (2) host innate/adaptive immune response – encompassing the complement cascade, leukocyte extravasation, ILK, IFN, chemokine, NF-kB, CD40, CCR3, and JAK-STAT signaling, (3) coagulation and thrombosis – involving renin-angiotensin, VEGF, endothelin, thrombin, angiopoietin, and erythropoietin signaling, (4) protein translation – including PI3K/AKT, p70S6K/mTOR, EIF2, and ERK/MAPK signaling, (5) cellular homeostasis – involving autophagy, sirtuin signaling, the TCA cycle, and glycolysis, and (6) fibrosis – such as hepatic and idiopathic pulmonary fibrosis. (**Figure 2c**). In addition, gene ontology enrichment analysis revealed functional terms related to SARS infection, cell cycle, protein phosphorylation, growth factor signaling, and neutrophil/platelet degranulation (**Figure S3f**). In parallel, kinase activity scores estimated from quantitative phospho-proteomic data showed that SARS-CoV-2 infection leads to activation of several kinases including members of the p38/MAPK pathway (MAPK1, MAPK13, ERK, ERBB3), CK1 (CSNK2A1), AKT/PI3K pathway (RSK, RPS6KA), Ca2+/calmodulin-dependent protein kinases (DAPK1, PRKAA2), PKC family (PRKCE, PRKCA, PRKCG), DNA damage response (DNA-PK, ATM) and stress response related proteins (SGK1). Kinases predicted to be downregulated included electrolyte homeostasis (WNK1), immune response (CAMK4, PRKCQ) and cytoskeletal rearrangement (EPHA2, ROCK) related regulators, among others (row2; **Figure 2g**). Similar pathway activation and kinase activity profiles were observed in PBMCs of severe/ICU COVID-19 patients and other *in vitro* models of SARS-CoV-2 infection ^42, 66–69^. SARS-CoV-2 infection also resulted in activation of other upstream regulators involved in coagulation/wound healing (EDN1, FN1) and cytokine production (CSF1, CCL2, IRS2, NCOA3) and suppression of factors involved in antiviral response (PTPRK, PPP2R1A) (**Figure 2h**). In line with previous reports ^51–56, 67^, network activity analysis also showed that SARS-CoV-2 infection mediated activation of translation (EIF2/4), hyperinflammation (IL1B, IL-18) and apoptosis/lymphopenia (CASP3/9) related pathways with concomitant suppression of anti-viral/anti-inflammatory response (NF-kB, IFN type-1, Jak-STAT) in A549-hACE2 cells (**Figure 4a & 4b**).

**Figure 3.**
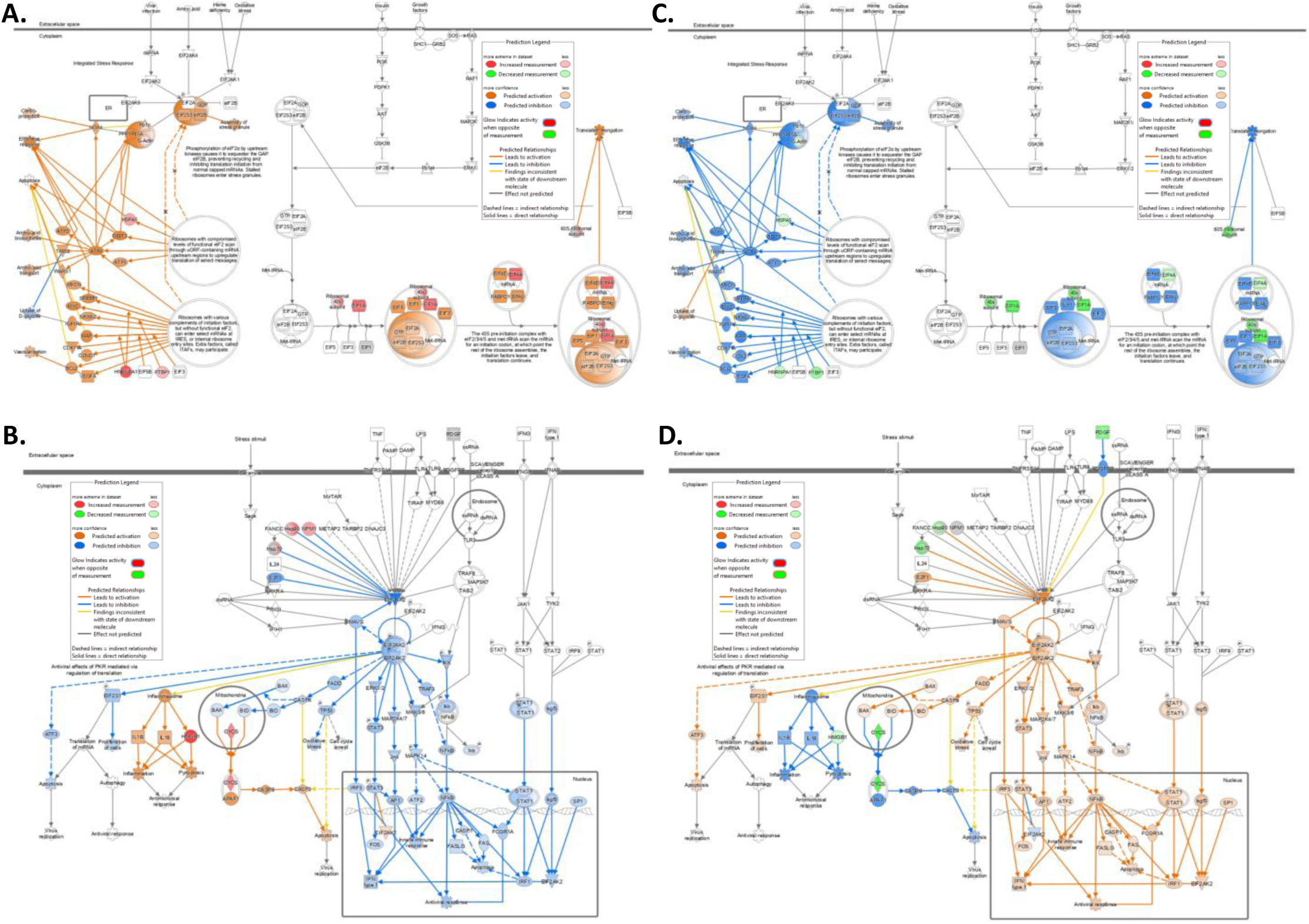
G9a inhibition reversed SARS-CoV-2 dysregulated host pathways related to translation and antiviral response. (**A-B**) Networks overlaid with gene expression changes following SARS-CoV-2 infection, compared to mock infected controls (i.e., SARS2 vs Ctrl), show activation of pathways related to translation (EIF2/4), hyperinflammation (IL1B, IL-18), and apoptosis/lymphopenia (CASP3/9) along with concomitant suppression of anti-viral/anti-inflammatory response (NF-kB, IFN type-1, Jak-STAT) in A549-hACE2 cells. By comparison, **(C-D)** UNC0642 treatment of SARS-CoV-2 infected cells (i.e., SARS2+UNC0642 vs SARS2) reversed these changes. Ingenuity Pathway Analysis was used to generate these networks overlaid with gene expression levels (RNA-Seq). Nodes colored red/green represent increased/decreased expression in our dataset, whereas orange/blue colored nodes and edges represent predicted activation/deactivation of said regulators based on our data. Yellow color refers to our identifications that are different from the literature, whereas grey color indicates unpredictable pathway activities. (See also **Figure S3, Supplemental Table S2**).

**Figure 4.**
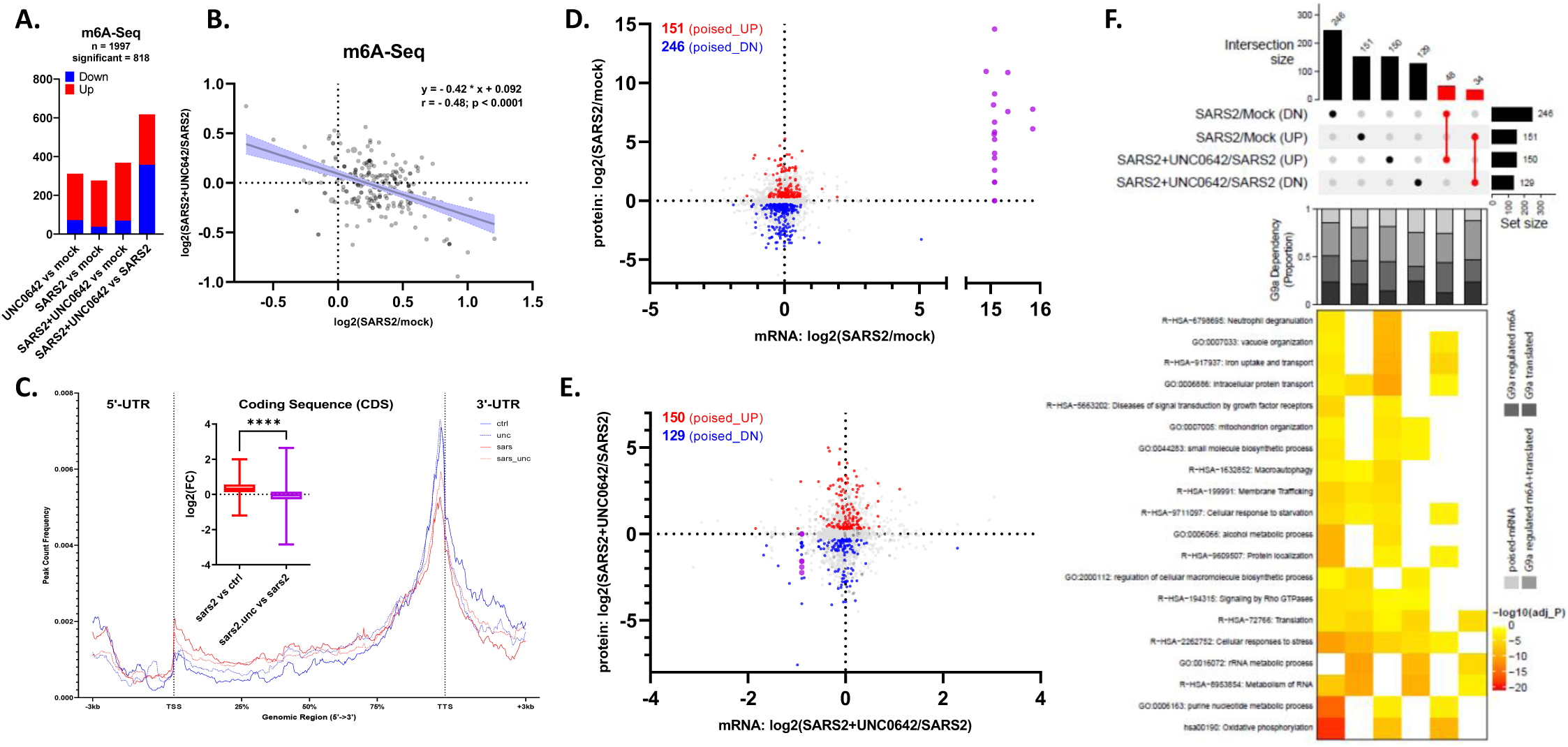
G9a and SARS-CoV-2 co-upregulate m^6^A modification of select mRNAs involved in proviral host response. (**A**) Number of transcripts with significant difference in m^6^A modification level, detected using m^6^A-Seq, following SARS-CoV-2 infection and/or UNC0642 treatment of A549-hACE2 cells **(B)** UNC0642 treatment reverses SARS-CoV-2-mediated changes in the host m^6^A methylome. Linear regression line (with 95% CI highlighted), slope of regression line (β) and Pearson correlation (r) are shown. **(C)** Distribution of the enriched m^6^A peaks, respectively, in control (blue line), UNC0642 treated (dotted blue line), SARS-CoV-2 infected (red line), and SARS-CoV-2 infected + UNC0642 treated (dotted red line) A549-hACE2 cells analyzed along the RNA segments. Each transcript was length normalized and ±3 kb from TSS/TTS are included. Boxplot shows an increase in m^6^A level following SARS-CoV-2 infection (red), compared with uninfected control, which is reversed upon UNC0642 treatment (magenta). Mann-Whitney test was used for statistical analysis (****p < 0.0001). **(D-E)** Scatter plots depicting transcriptomic and proteomic changes following SARS-CoV-2 infection (D) and UNC0642 treatment (E) of infected A549-hACE2 cells compared to respective controls. Poised mRNAs with statistically significant increases (red) or decreases (blue) in protein expression are highlighted. Number of poised mRNAs (i.e., genes showing differential protein expression without detectable change at the transcript level) is shown (top left). SARS-CoV-2 encoded genes/proteins (showing increases upon SARS2 infection and decreases upon UNC0642 treatment) are highlighted by purple dots. **(F)** On top is upset plot showing number and overlap among poised mRNAs identified in (D) and (E). Poised mRNAs dysregulated following SARS-CoV-2 infection whose expression was reversed following UNC0642 treatment are highlighted in red. Stacked bar plot in the middle shows proportion of identified poised mRNAs whose m^6^A, translation, or both m^6^A and translation is regulated by G9a/UNC0642. Pathway enrichment for indicated gene sets is shown at bottom. (See also **Figures S4 & S5, Supplemental Table S3**)

We also evaluated the contribution of RNA/protein abundance to changes at either phosphorylation or secretion level. Consistent with reports of increased cytokine secretion^70, 71^ and host phosphorylation landscape rearrangement^67^, for nearly all cases of a significantly changed phosphorylation site and/or secreted protein, we did not observe corresponding significant changes in RNA/protein abundance **(Figure S3g)**. These results suggested that, instead of transcriptional regulation, post-translational (i.e., phosphorylation or secretion) regulation is the primary host response to SARS-CoV-2 infection, which would influence protein abundance, at least during the timescale of infection of our study.

### G9a inhibition rescues SARS-CoV-2 hijacked pathways

As evidenced by negative overall correlations (RNA-Seq = –0.53; proteome = –0.48; phosphoproteome = –0.47; secretome = –0.42), pharmacologic inhibition of G9a led to reversal of SARS-CoV-2-mediated changes in abundance (RNA-Seq, proteome), secretion (secretome), and phosphorylation (phosphoproteome) of various viral/host transcripts and proteins in A549-hACE2 cells (**Figure 2b**). We observed similar reversal in the activity of SARS-CoV-2 dysregulated pathways following G9a inhibition, an effect that was absent in UNC0642-treated mock-infected controls, thereby further associating G9a activity with viral replication and the SARS-CoV-2 dysregulated host response (**Figure 2c**). Specifically, abundance, secretion, and/or phosphorylation patterns of 351 host or viral entities dysregulated following SARS-CoV-2 were reversed upon UNC0642 treatment. Most of these entities showing SARS-CoV-2-indcued, UNC0642-reversed patterns in “Cluster-1” were related to coronavirus pathogenesis, including viral transcripts and proteins, host receptors for virus entry (notably, ACE2), fibrosis markers, cytokines, and coagulation or angiogenesis-related proteins (**Figure 2d**). Nearly all members of Cluster-1 were host interactors of various SARS-CoV-1/2– and MERS-CoV-encoded proteins, further highlighting G9a’s function in promoting expression/phosphorylation of host factors necessary for SARS-CoV-2 infection/replication (**Figure 2e**). Overall, these SARS-CoV-2 promoted, UNC0642-reversed, host-interactors were involved in complexes or pathways related to immune response (e.g., antiviral/stress response, neutrophil degranulation, cytokine signaling), coagulation or angiogenesis (e.g., platelet aggregation, VEGF signaling), translation (e.g., ribosome biogenesis), energy metabolism (e.g., carbohydrate, carbon, small molecule synthesis), and cell cycle (**Figure 2f & S3h**).

In parallel, we evaluated the effect of G9a inhibition by UNC0642 on the host response of mock-infected A549-hACE2 cells. Although UNC0642 treatment altered expression, secretion and/or phosphorylation of certain host proteins (**Figure S3d**), overall, there was negligible effect on global the phosphoproteomic landscape. Thus, G9a had little effect on host phosphorylation-dependent signaling under noninfected circumstances (i.e., Row 1 in **Figure 2g**). More importantly, in line with global reversal of SARS-CoV-2-mediated changes in abundance and phosphorylation of host and viral proteins, UNC0642 treatment reduced or reversed the kinase activity profile of SARS2 infection (Row 3 in **Figure 2g**). Coincidently, therapeutic targeting of several G9a-regulated kinases that we identified inhibited SARS-CoV-2 infection and replication in vitro^42, 66–68^. These kinases are functionally associated with growth factor signaling (e.g., MAPK, PI3K/Akt, TGF-b), protein translation (e.g., mTOR, p38/MAPK), cell cycle (e.g., CDKs), and cytoskeletal rearrangement (e.g., CSNK2A1/2). Similarly, G9a inhibition did not affect the activities of regulators related to coagulation/wound healing, cytokine production, and antiviral response in mock-infected cells but, in contrast, led to stark reversal in activation of these regulators in SARS-CoV-2-infected A549-hACE cells (**Figure 2h**). Network activity analysis also showed G9a inhibition-reduced activation of the pathways related to translation (e.g., EIF2/4), hyperinflammation (e.g., IL1B, IL-18), and apoptosis/lymphopenia (e.g., CASP3/9) along with concomitant increases in the pathways related to anti-viral/anti-inflammatory response (e.g., NF-kB, IFN type-1, Jak-STAT) (**Figures 4c & 4d**). On a pathway scale, these data explain G9a’s function in mediating SARS-CoV-2 rewiring of host signaling to promote viral replication and dysregulated immunity.

Yaron et al. implicated phosphorylation of SARS-CoV-2 proteins in viral replication/life cycle and pathogenesis ^72^. Correspondingly, we identified six phosphorylation sites on SARS-CoV-2 encoded proteins, including Orf9b (Ser50), M (Ser213) and N (Ser78, Ser176, Ser412, Ser413) (**Figure 2b, Supplemental Table S2c**). These sites are conserved among coronaviruses, indicative of functional constraint. Top kinase families predicted to phosphorylate these sites include casein kinase II (CK2), glycogen synthase kinase (GSK-3), ribosomal s6 kinase (RSK), and protein kinase C (PKC), suggesting that these kinases may contribute to regulation of viral replication^67, 72, 73^. This idea is in line with top upstream kinases predicted to be regulated by UNC0642 in our dataset to reverse phospho-proteomic landscape of SARS2 infection. More importantly, UNC0642 treatment of SARS2-infected cells resulted in marked decrease in phosphorylation at four viral sites, including ones on N (Ser78, Ser176, Ser412) and Orf9b (Ser50), compared to SARS2 infected cells alone (**Figure 2b**). Mechanistic details and potential functionality of these G9a regulated sites are unknown and warrant further investigation.

In sum, our multiomics correlation analysis revealed that UNC0642 treatment, which had minimal effect in noninfected cells, broadly rescued the widespread SARS-CoV-2-hijacked signaling pathways. Mechanistically, G9a activity regulates the SARS-CoV-2-mediated activities of translation-regulatory kinases such as RSK, supporting the translation-regulatory function of G9a in SARS-CoV-2 replication/life cycle and the dysregulated host response.

### G9a promotes SARS-CoV-2/m^6^A-coregulated translation for COVID-19 pathogenesis

METTL3 (m^6^A writer) stabilizes mRNA transcripts by introducing m^6^A to promote translation^74, 75^. In addition, SARS-CoV-2 infection increases expression of m^6^A regulators, including METTL3, and induces COVID-19 characteristic m^6^A landscapes in both virus and the host cells or patients, wherein the m^6^A modification was implicated in viral replication and dysregulation of host immune response^33–40, 76–78^. Similarly, we showed that, in ET macrophages, G9a and METTL3 co-upregulated translation of certain m^6^A-modified mRNAs associated with sepsis and ARDS. Depletion of G9a or METTL3 restored T cell function and reduced macrophage activation syndrome ^79^. Further, in line with reports of increased expression of RBM15 and WTAP in COVID-19 patient PBMCs ^33^, we observed increased expression of METTL3 complex components (RBM15, WTAP) and m^6^A readers (TRA2A) upon SARS-CoV-2 infection, which was suppressed by UNC0642 treatment in SARS-CoV-2 infected cells (**Figure S3i**). In correlation with UNC0642-suppressed SARS-CoV-2 replication (**Figure 1e & 2B**), our results highlighted a triggering possibility that G9a is the upstream regulator of SARS-CoV-2 hijacked, m^6^A-mediated translation of proteins associated with COVID-19 pathogenesis. Thus, targeting G9a represents both a virus– and a host-directed mechanism of COVID-19 therapeutic action.

To investigate exactly how active G9a assists SARS-CoV-2 to hijack the host translation pathways, we characterized the function of the ChaC-identified interaction between G9a and METTL3 in SARS-CoV-2 induced pathogenesis. Specifically, we examined whether and how G9a activity would influence the distribution of m^6^A on cellular transcripts during infection. Accordingly, using MeRIP-Seq, we identified m^6^A-modified transcripts that had SARS-CoV-2 induced changes and which were reversed by UNC0642 treatment in infected A549-hACE2 cells. Like other omics datasets, we observed a clear principal component separation between mock and infected cells, and UNC0642 treatment resulted in distinct host m^6^A landscape (**Figure S4a**). Nearly two thousand m^6^A-modified transcripts were identified with 818 transcripts having differential m^6^A abundance following infection and/or UNC0642 treatment (**Figure 4a; Supplemental Table S3a**). Coincident with elevated m^6^A regulator expression in A549-hACE2 cells following SARS-CoV-2 infection (**Figure S3i**), and previous reports for A549-hACE2 cells and COVID-19 patient PBMCs ^40, 78, 80^, we detected an overall increase in m^6^A abundance following infection, compared with uninfected controls; this effect was reversed upon UNC0642 treatment as evidenced by the overall negative correlation of –0.48 (**Figure 4b-4c**). Specifically, SARS-CoV-2 infection led to increased m^6^A modification in the 5’-UTR and coding sequence (CDS) regions along with a concomitant decrease in m^6^A abundance around the 3′-UTR of host transcripts. However, UNC0642 treatment reversed these SARS-CoV-2-mediated changes in CDS m^6^A levels (decrease) and 3’-UTR (increase) regions, and led to decreased m^6^A in the 5’-UTR regions in general (**Figure 4c**). Most SARS-CoV-2 dysregulated, UNC0642-reversed, m^6^A-modified transcripts were involved in pathways related to type I/II interferon regulated genes involved in immunity (viral infection, neutrophil degranulation), cell cycle (mitosis, G2/M transition), translation, RNA splicing/processing, cytoskeletal rearrangement, and energy metabolism (carbohydrates, glucose, glycolysis, steroids) (**Figure S4b-S4c**). Interestingly, most transcripts with differential m^6^A did not exhibit corresponding changes in mRNA expression (**Figure S4d**); thus, increased m^6^A following SARS-CoV-2 infection was not due to changes at the transcriptional level but, instead, due to a post-transcriptional host response to virus infection.

### G9a directs translation of viral and host proteins for SARS-CoV-2 replication and dysregulated host response

Following virus entry into host cells, viral mRNAs are ‘poised’ to rapidly undergo translation and produce virus replication proteins and trigger translation of specific mRNAs for the host response. However, little is known about how these poised mRNAs are regulated during SARS-CoV-2 pathogenesis. To further elucidate the mechanism of G9a-mediated translation in SARS-CoV-2 replication and associated pathogenesis, we compared transcriptomic (RNA-Seq) and LFQ proteomic datasets from the A549-hACE2 cells with versus without SARS-CoV-2 infection or with and without treatment of G9a inhibitors. These multi-omics comparisons identified G9a-regulated, poised mRNAs, i.e., genes that showed changes in protein expression without any significant change in corresponding mRNA levels following SARS-CoV-2 infection. As expected, we detected 397 (151 upregulated; 246 downregulated) and 279 (150 upregulated; 129 downregulated) differentially expressed poised mRNAs following SARS-CoV-2 infection with versus without UNC0642 treatment (**Figure 4d-4e; Supplemental Table S3b**). SARS-CoV-2 infection reshaped host signaling to produce more protein from viral infection, translation/rRNA-biogenesis and cell cycle related mRNAs, whereas infection reduced translation of host immunity, stress response, and energy metabolism related genes (**Figure 4f**; columns 1 to 4). More importantly, these SARS2 mediated changes in expression of host poised mRNAs were largely mitigated (120 total; 48 down/up; 34 up/down) following UNC0642 treatment of infected cells (**Figure 4f**; columns 5-6). Lastly, the majority of SARS-CoV-2 regulated poised mRNAs whose expression was reversed by UNC0642 were also identified as ‘G9a-translated’ proteins that carried G9a/METTL3-corregulated m^6^A modification in ET macrophages (**Figure 4f**). These results underpinned the crucial activity of G9a in SARS-CoV-2 mediated hijacking of host translational machinery to increase virus production and evade host cell immune responses. Together, we showed that G9a upregulated m^6^A of select host mRNAs following SARS-CoV-2 infection to promote translation of transcripts functionally related to immune response, rRNA/ribosome, and viral entry/egress.

### SARS-CoV-2-induced, G9a-mediated translation contributes to COVID-19 complications

To ascertain the clinicopathologic relevance of the G9a translational mechanism of COVID-19 pathogenesis, we conducted transcriptomic (RNA-Seq), epi-transcriptomic (m^6^A/RIP-Seq), and LFQ proteomic experiments to identify G9a-regulated, poised mRNAs from *ex vivo* culture of COVID-19 patient derived PBMCs. Briefly, akin to the results from A549-hACE2 cells, we identified 292 differentially expressed proteins that were coded by poised mRNAs, most without a corresponding change in transcript level, following UNC0642 treatment of patient PBMCs **(Figure 5a; Supplemental Table S4a)**. Several proteins were host interactors of various SARS-CoV1/2– and MERS-CoV-encoded proteins and, partially by m^6^A modification of transcripts, their turnover/translation was regulated in a G9a-dependent manner in ET macrophages **(Figure 5a)**. Overall, G9a regulates protein expression/abundance of transcripts related to translation, immunity/infection, lipid metabolism, DDR, cellular energetics, ubiquitination, and unfolded protein response in patient PBMCs (**Figure 5b)**. Like A549-hACE2 cells, UNC0642 treatment of COVID19 patient PBMCs also reversed SARS-CoV-2 regulated m^6^A modification in the CDS (decrease) and 3’-UTR (increase) regions of host transcripts. However, unlike A549-hACE2 cells, overall m^6^A (particularly in the promotor/5’-UTR region) increased following UNC0642 treatment (**Figure S5a**), possibly owing to cell-typic differences in m^6^A regulator expression. Overall, most m^6^A modified transcripts in control (DMSO) patient PBMCs belonged to type I/II interferon regulated genes (**Figure S5b**) involved in adaptive immune response pathways, translation, infection/endocytosis, blood coagulation, and antibiotic response, a pattern largely absent from UNC0642-treated patient PBMCs (**Figure S5c**).

**Figure 5:**
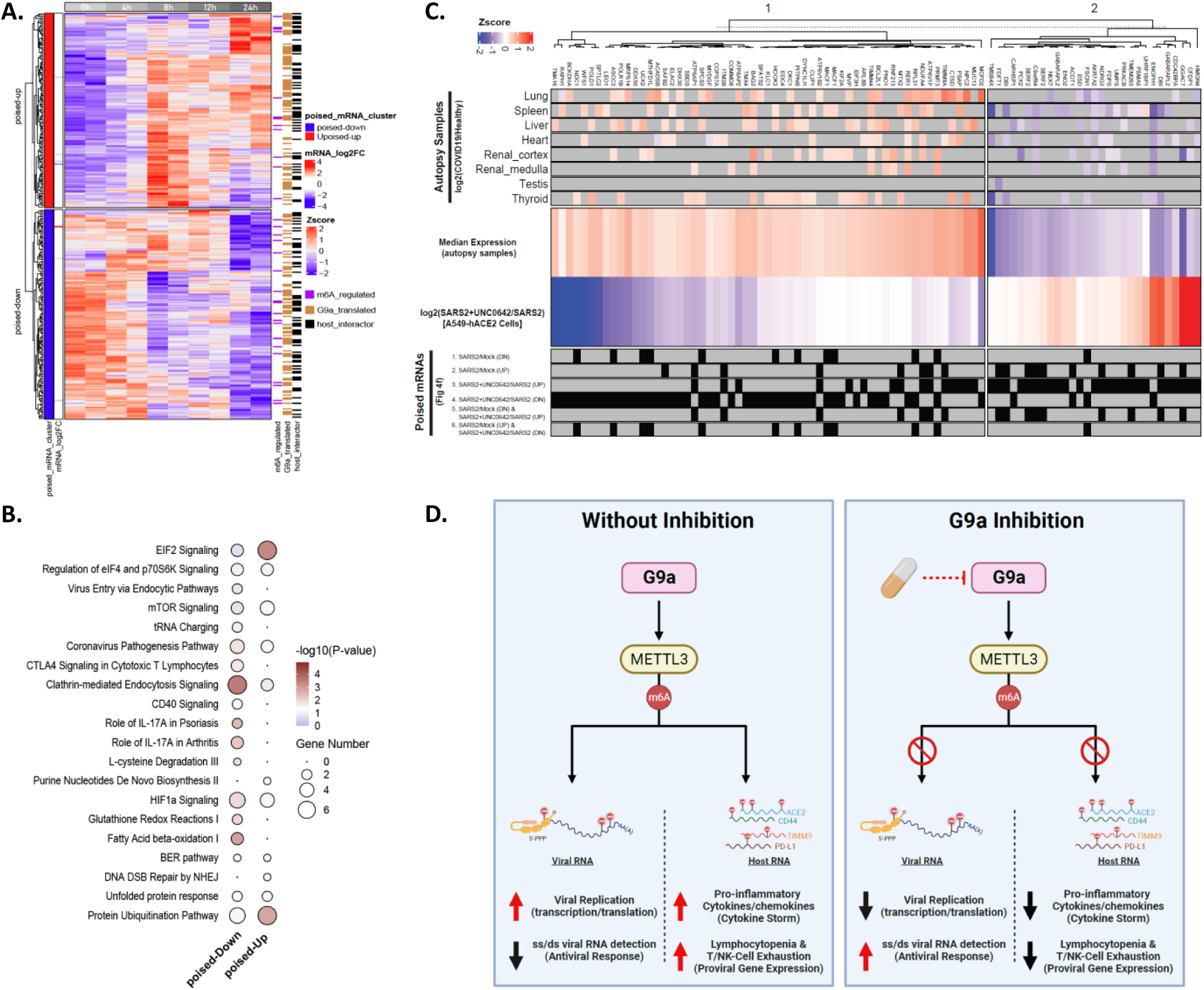
G9a regulates poised mRNA expression in COVID-19 patient PBMCs and autopsy samples. (**A**) Heatmap summarizing transcriptomic (RNA-Seq), epi-transcriptomic (meRIP-Seq), and proteomic (LFQ-MS) effects of UNC0642 inhibition in COVID-19 patient PBMCs. Briefly, *ex-vivo* cultures of patient PBMCs were treated with 1 μM UNC0642/DMSO and samples collected at indicated time points for LFQ-MS; RNA-Seq and meRIP-Seq were performed at 24 h. In line with A549-hACE2 results, most UNC0642 regulated genes showed differential protein expression without detectable change at transcript level (i.e., poised mRNAs). Annotations on right show that most of these genes are translated in a G9a-dependent manner in ET macrophages, with a subset carrying G9a regulated m^6^A modification, and are host interactors of SARS-CoV– 1/2– and MERS-CoV-encoded proteins. **(B)** Pathway enrichment analysis for G9a regulated poised mRNAs shown in (A). All terms with significant over-representation (adjusted P < 0.05) are kept, and redundant terms are removed. Dot size corresponds to the number of dysregulated genes/proteins in said term. **(C)** Heatmap showing that several proteins with multi-organ dysregulation in autopsy samples from COVID-19 patients are, in fact, encoded by SARS-CoV-2/G9a-coregulated poised mRNAs, and their expression was reversed upon UNC0642 treatment of SARS-CoV-2 infected A549-hACE2 cells. Rows 1-8: log2(COVID-19/healthy) ratio for indicated organs; Rows 9: Median expression of indicated proteins in autopsy samples. Row 10: log2(SARS2+UNC0642/SARS2) ratio in A549-hACE2. Rows 11-16: indicate whether said protein is encoded by poised mRNAs identified in earlier (Fig. 4f) comparisons. **(D)** Possible mechanisms of G9a action. Briefly, SARS-CoV-2 infection uses G9a-METTL3-m6A axis to promote m^6^A modification on (1) host transcripts to promote expression of lymphopenia, T/NK cell exhaustion and hyperinflammation related proteins, and (2) SARS-CoV-2 genome to promote viral transcription/translation and evade RIG-1 dependent sensing and activation of innate immune response and ensuing IFN-β/type-I IFN antiviral response. We show that G9a inhibition reverses SARS-CoV-2 mediated rewiring of m^6^A epi-transcriptome to hinder SARS-CoV-2 replication, suppress expression of pro-inflammatory cytokines, and reduce expression of markers associated with T/NK-cell exhaustion, including PD-L1 and other proteins involved in lymphopenia. (See also **Figure S5, Supplemental Table S4**)

Notably, several of the virus/G9a-dysregulated and inhibitor-reversed entities identified earlier (**Figure 2d**) are known markers of Long COVID or Post-Acute Sequelae of SARS-CoV-2 infection^81^, blood/plasma markers of severe COVID19^82, 83^, and are dysregulated in autopsy samples ^81^ (**Figure S5e**). Thus, from our A549-hACE2 global proteomics (LFQ+TMT) datasets, we assessed clinical data^29^ to identify 88 SARS-dysregulated/G9a-reversed proteins that showed similar aberrant expression in multi-organ proteomics data from autopsy samples (**Figure 4c**), with nearly all transcripts carrying G9a regulated m^6^A modification in ET macrophages, A549-hACE2 cells, and/or COVID19 patient PBMCs **(Supplemental Table S4b)**. As expected, these G9a regulated patient-specific proteins were involved in immunity (neutrophil degranulation/VEGF signaling/stress response), energy metabolism, and cellular transport/localization pathways (**Figure S5d**). Taken together, we show that G9a regulates translation of certain poised mRNAs in COVID-19 patient PBMCs and autopsy samples following SARS-CoV-2 infection, at least partly by the m^6^A modification pathway, further establishing the clinicopathological relevance of our multi-omics results. More significantly, these COVID-19 proteomic and phosphoproteomic landscapes can be reversed by inhibitor treatment, indicating possible clinical efficacy of G9a inhibitors for COVID-19 therapy or treatment of COVID-19 sequalae.

### Pharmacological intervention of G9a-associating translation machinery suppressed SARS-CoV-2 pathogenesis

Because EZH2 may be a key factor in G9a-mediated translational regulation of COVID-19 pathogenesis, we inhibited EZH2 with UNC1999 and measured proteome and phosphoproteome changes in SARS-CoV-2 infected A549-hACE2 cells. UNC1999 treatment resulted in a distinct proteomic landscape compared with controls (**Figure 6a-6b; Supplemental Table S5a-S5b**) with good reproducibility between and across replicates (**Figure 6c**). From a systems view, UNC1999 treatment led to significant reduction in expression of SARS-CoV-2 encoded proteins along with overall reversal in SARS-CoV-2 induced changes in the host proteome (correlation = –0.349; **Figure 6d**) and phosphoproteome (correlation –0.281; **Figure 6e**). Notably, similar to effects of G9a inhibition, UNC1999 treatment reversed SARS-CoV-2 induced changes primarily at the phosphoproteomic level as opposed to changes in protein abundance in SARS-CoV-2 infected cells (**Figure 6d-6e**). Most of these UNC1999-affected phosphoproteins are functionally associated with coronavirus replication, virus entry/egress, immune response, and cholesterol signaling (**Figure 6f**). Specifically, on the basis of quantitative phosphoproteomic identification of SARS-CoV-2-induced proteins whose phosphorylation was –reversed by UNC1999, we found that SARS-CoV-2 infection activated multiple kinase-mediated signaling pathways, including MAP/ERK pathway (MAPK1, MAPK13, ERK), RTK family (EGFR, PDGFR), cell cycle (CDK1/6 CCNB1, CDC2), Casein kinase (CSNK2A1/2), AKT/PI3K pathway (MTORC1, RPS6KA1/B1, AKT3), Ca2+/calmodulin-dependent protein kinases (DAPK1, PRKAA2), PKC family (PRKCA, PRKCE, PRKCG) and DDR (ATM, PLK1) related kinases. In parallel, decreased activity was observed for specific kinases related to cell cycle (CDKN1B) and cytoskeleton regulator (ROCK2) (**Figure 6g**). Overall, the abundance and/or phosphorylation pattern of 235 entities dysregulated by SARS-CoV-2 infection were reversed upon UNC1999 treatment, including several coronavirus pathogenesis related entities, e.g., viral transcripts/proteins, host receptors (ACE2), fibrosis markers, cytokines, coagulation/angiogenesis-related proteins, and transcription factors (**Figure 6h**). Most of these proteins were host interactors of SARS-CoV-1/2– MERS-CoV-encoded proteins, further highlighting Ezh2’s function in promoting expression and/or phosphorylation of host factors necessary for SARS-CoV-2 infection and replication (**Figure 6i**). Together, these SARS-CoV2-induced/UNC1999-reversed host-interactors were involved in antiviral/stress response, cell cycle, RNA metabolism, protein transport, and DDR related pathways (**Figure 6j**).

**Figure 6.**
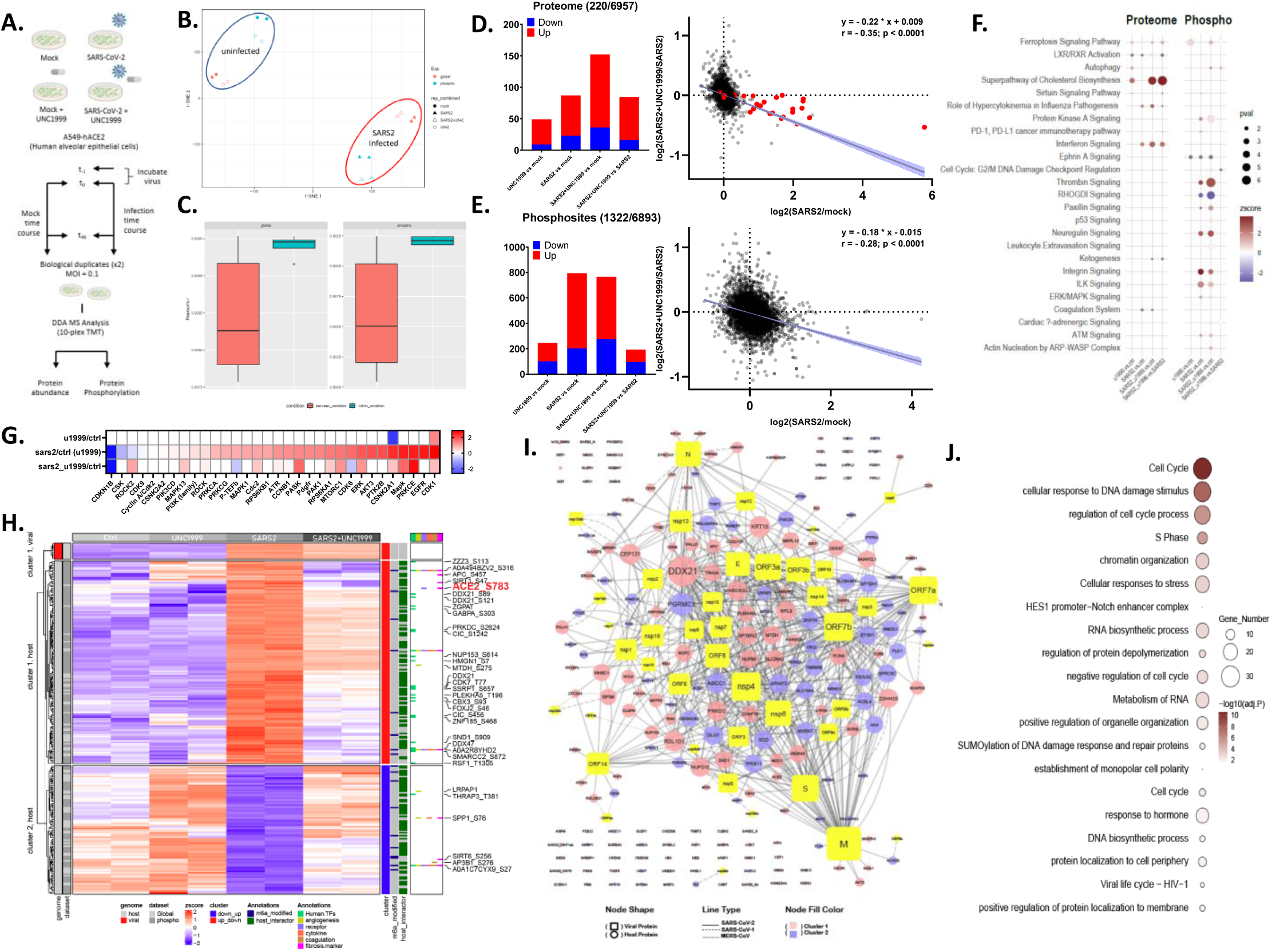
Ezh2 inhibition reverses proteomic landscape of SARS-CoV-2 infection. (**A**) Schematic overview of SARS-CoV-2 infection and Ezh2 inhibitor treatment of A549-hACE2 cells. Briefly, cells were infected with SARS-CoV-2 (MOI = 0.1). After 1 h of virus uptake, media were switched, and cells harvested 48 hpi with/without UNC1999 treatment. Proteins were digested and 10-plex TMT-labeled followed by DDA-based MS to detect global changes in protein abundance and phosphorylation. All experiments were performed in biological duplicate. **(B)** The t-distributed stochastic neighbor embedding (t-SNE) visualization of significantly dysregulated proteins in tested conditions (shape) and datasets (color). **(C)** Correlation between replicates within a biological condition (red) and across biological conditions (green). Boxplots depict median (horizonal lines), interquartile range (boxes), maximum and minimum values (vertical lines), and outliers (solid circles). **(D)** Bar plots depicting number of dysregulated proteins (red = up; blue = down) for indicated comparisons in UNC1999-treated proteomic dataset from A549-hACE2 cells. Total number of identified (n) and dysregulated (significant) entities is mentioned on top. Scatter plot on right shows that UNC1999 treatment reverses SARS-CoV-2 infection mediated changes to host (black) and viral (red) proteins in A549-hACE2 cells. Linear regression (with 95% CI), slope of regression line (β) and Pearson correlation (r) are overlaid. **(E)** Bar plot shows number of dysregulated phospho-sites (red = up; blue = down) for indicated comparisons in A549-hACE2 cells. Total number of identified (n) and dysregulated (significant) entities is mentioned on top. The scatter plot on right shows that UNC1999 treatment reverses SARS-CoV-2 infection mediated changes host phospho-proteome. Linear regression (with 95% CI), slope of regression line (β) and Pearson correlation (r) are overlaid. **(F)** Plot summarizing pathway activation scores estimated using differentially regulated entities in global/phospho-proteomic datasets following SARS-CoV-2 infection and/or UNC1999 treatment of A549-hACE2 cells. Each column corresponds to indicated pairwise comparison. Red represents pathway activation, and blue represents pathway inhibition. The dot size corresponds to adjusted p-values. Redundant terms were removed. **(G)** Kinase activity scores (– log10(P) < 0.05) estimated based on differentially regulated phospho-sites identified in A549-hACE2 cells following SARS-CoV-2/mock infection with or without UNC1999 treatment. Rows represent indicated pairwise comparison, and columns correspond to individual kinase/family (red = activation, blue = deactivation). **(H)** Heatmap of SARS-CoV-2 dysregulated, UNC1999 reversed, proteins/phospho-sites in A549-hACE2 cells. Viral and host entities are clustered separately and further subdivided into two groups (cluster1 = up following infection and down upon UNC1999 treatment; cluster2 = down upon infection and up upon UNC1999 treatment). Annotations on right highlight m^6^A-modified transcripts (identified by MeRIP-Seq) and host interactors of SARS-CoV-1/2– and MERS-CoV-encoded proteins. Lastly, six functional clusters of viral receptors and proteases, transcription factors, cytokines/chemokines, coagulation system-related proteins, angiogenesis-associated proteins, and fibrosis markers are included with their names highlighted. **(I)** Virus-host interactome map showing that most of the proteins identified in (H) are host interactors of SARS-CoV-2 (solid line), SARS-CoV-1-(dashed line) or MERS-CoV-(dotted line) encoded proteins. Viral proteins are depicted by yellow rectangles, and host interactors are represented by circles. Size of each node represents connectivity, and node fill color (cluster1 = pink; cluster2 = blue) represents coronavirus-dysregulated/Ezh2-reversed proteins. **(J)** Gene Ontology enrichment analysis for proteins shown in (H-I). All terms with significant over-representation (adjusted P < 0.05) are kept, and redundant terms are removed. Dot size represents the number of dysregulated proteins belonging to said term. (See also **Figure S6, Supplemental Table S5**).

Pharmacological inhibitors of Ezh2 are approved for treatment of epithelioid sarcoma and follicular lymphom;^63^ therefore, the inhibitors could be repurposed for COVID-19 therapy. Thus, we investigated the effect of Tazemetostat, the FDA-approved inhibitor of Ezh2, on coronavirus-related proteins in clinical setting, i.e., *ex vivo* culture of PBMCs from two severe COVID-19 patients. Out of 4221 proteins quantified by LFQ-MS, 420 proteins (305 down, 115 up) were differentially regulated following treatment (**Figure S6a; Supplemental Table S5c**). Strikingly, 193 dysregulated proteins (157 down, 36 up) were host interactors of SARS-CoV-1/2– and MERS-encoded proteins, clearly demonstrating that these Tazemetostat-downregulated proteins are required for efficient SARS-CoV-2 replication or infection. Overall, Tazemetostat-dysregulated proteins were involved in viral-entry (i.e., coronavirus pathogenesis, endocytosis, phagocytosis), translation (EIF2, mTOR, eIF4 and p70S6K), immune response (interleukin/cytokine, neutrophil, macrophage, inflammation, NF-kB, apoptosis, autophagy) and cellular metabolism (nucleotide, cholesterol, amino-acid) related signaling (**Figure S6b**). Several proteins in these pathways are among the top hits/host factors necessary for coronavirus pathogenesis, characterized by multiple genome-wide siRNA/CRISPR studies.^21, 62, 84–87^ For example, Tazemetostat suppressed the expression of 12 host factors known to hinder SARS-CoV-2 infection, including CSTL/VPS34 (spike cleavage and membrane fusion), PIK3C3 (endosome maturation) and ERMP1 (ER/Golgi-trafficking) (**Supplemental Table S5c**). Systematic analysis of disease-related signaling cascades showed a strong suppression of viral/bacterial infection and immune system activation (**Figure S6c**), further corroborated by UNC1999-mediated suppression of translation (EIF2/4) and inflammation (NF-kB, IL-6, IL-8) related network activity (**Figure S6d-S6e**). Together, Ezh2 inhibition compromised SARS-CoV-2 mediated changes to host proteome and phosphoproteome to hinder SARS-CoV-2 replication in A549-hACE2 cells and, more importantly, suppressed (1) host interactors of SARS-CoV-2 encoded proteins, (2) host factors required for efficient SARS-CoV-2 infection/replication, and (3) critical pathways involved in coronavirus pathogenesis in COVID-19 patient PBMCs. Combined results indicated that EZH2 is a key component of the G9a translational regulatory machinery whereby G9a and Ezh2 cooperatively promote SARS-CoV-2 pathogenesis primarily at the translational and/or post-translational level.

## DISCUSSION

We investigated how SARS-CoV-2 hijacks the host response mechanism to produce specific proteins that are the primary executers of COVID-19 pathogenesis. In line with the fact that dysregulated translation of specific proteins directly contributes to disease pathogenesis, we discovered that, via interactions with translation regulators such as METTL3 and Ezh2, the histone methyltransferase G9a broadly regulates widespread SARS-CoV-2-hijacked translation pathways^30^ associated with viral replication, hyperinflammation, T cell exhaustion, and suppressed host antiviral response (**Figure 5d**, left panel). In parallel, we revealed the mechanism of inhibitor action for both virus– and host-directed therapeutics of COVID-19, whereby G9a/Ezh2 inhibitors reverse SARS-CoV-2 dysregulated translation processes (**Figure 5d**, right panel). Given higher resistance barrier, broader activity against coronavirus strains/species, and potential synergy with other direct-acting antiviral drugs^88, 89^, our approach of targeting G9a-regulated mechanisms of COVID-19 pathogenesis represents broad-spectrum, precision therapeutics to counter any emerging coronavirus variant and to prevent COVID-19 sequelae.

Evidently, m^6^A modification is the most prevalent hallmark of SARS-CoV-2-hijacked translation pathways. Increased viral m^6^A modification stabilized SARS-CoV-2 transcripts^34^ to promote replication and translation of SARS-CoV-2 genome^33–35^ and to evade RIG-1-dependent sensing and activation of the innate immune response^76^. Simultaneously, SARS-CoV-2 infection rewired the host m^6^A epitranscriptome to promote programmed cell death in lymphocytes ^33^ and upregulate expression of pro-viral and inflammatory response genes^33, 35, 77, 78^. As an m^6^A writer, METTL3 interacts with SARS-CoV-2 RNA-dependent RNA polymerase (RdRp) in the host to facilitate SARS-CoV-2 RNA m^6^A modification and viral replication^35^. Similarly, an elevated level of METTL3 and m^6^A readers, such as RBM15B, IGF2BP1, hnRNPA1, coupled with a decrease in m^6^A demethylase (FTO, ALKBH5) expression was reported in SARS-CoV-2 infected cell lines^33–36^ and clinical isolates from severe COVID-19 patients^37–40^. Correspondingly, we identified by ChaC most of m^6^A regulators (e.g., writers or readers) as the COVID-19 phenotypic interactors and showed that G9a inhibition reversed the SARS-CoV-2-induced m^6^A landscape (**Figure 4b-4c**); thus, via the METTL3-m^6^A axis, G9a regulates select SARS-CoV-2-hijacked translation pathways. For example, in correlation with the observation that compromised antigen presentation by METTL3^hi^CD155^hi^ macrophage cells diminishes antiviral T cell response against SARS-CoV-2 antigens in COVID-19 patients with coronary artery disease^90^, we found that G9a inhibition reversed the level of select m^6^A mRNAs associated with macrophage proliferation, T-cell dysfunction, and dysregulation of blood coagulation. Presently, in the absence of good clinical/pre-clinical molecules targeting m^6^A regulators (METTL3), our results demonstrate that inhibition of G9a, the upstream regulator of SARS-CoV-2 induced m^6^A epitranscriptome, represents a host mechanism-directed therapeutic approach to combat SARS-CoV-2 infection.

Multiomics-correlated analysis identified poised mRNAs whose translation was triggered by SARS-CoV-2 infection in a G9a activity-dependent manner. As a result, G9a translates these poised mRNAs into proteins that promote SARS-CoV-2 replication, including ACE2 binding/entry and pathogenesis. Further, phosphoproteomic data showed that G9a regulated host kinase signaling, indicating that most regulation of SARS-CoV-2 pathogenesis occurs by post-translational modification (phosphorylation or secretion) as opposed to regulating protein abundance.

Pathogenic influenza and β-coronaviruses (SARS-CoV-1/2 and MERS-CoV) suppress transcription of IFN-regulated genes^91–93^ by multiple mechanisms that include increased repressive marks (H3K9me and H3K27me) and histone mimicry employed by SARS-CoV-2^57, 58^. Interestingly, Hu et al. reported that EZH2 interacts with METTL3 ^59^ and Yi et al. described a PRC2-independent function for EZH2 in IRES-dependent translation^60^, raising the possibility that EZH2 might be a key component of the G9a-associated translational regulatory machinery that promotes COVID-19 pathogenesis. In this study, we showed that multiple members of G9a, EZH2, and METTL3 complexes were upregulated in nasopharyngeal swabs and/or autopsy samples from severe COVID-19 patients (**Figure S1a-S1b**). G9a and Ezh2 interacted with the same translational regulators in ET macrophage and COVID-19 patient PBMC cells (**Figure 1b-1c**) to help G9a promote turnover of host interactors of SARS-CoV-1/2– and MERS-encoded proteins and other pro-viral host factors (**Figure 1d**). Drugs targeting G9a and Ezh2 were potent inhibitors of SARS-CoV-2 replication (**Figure 1e**) and reversed multi-omic effects of SARS-CoV-2 infection in A549-hACE2 cells and COVID-19 patient PBMCs (**Figures 2-6, S3-S6**). We identified a mechanistic axis by which G9a remodels viral and host m6A epi-transcriptome following SARS-CoV-2 infection (**Figure 4-5**) to promote viral replication and infection by upregulating expression of various viral receptors (including ACE2), transcription factors, cytokines (and their receptors), coagulation/angiogenesis related proteins, and fibrosis markers in A549-hACE2 & COVID-19 patient PBMCs (**Figure 2**) with said proteins showing similar dysregulation pattern in multiorgan autopsy samples taken from deceased COVID-19 patients (**Figures 5**). Together, we showed that G9a inhibition suppressed SARS-CoV-2 replication while reducing inflammatory cytokine and proviral gene expression and enhancing antiviral response (**Figure 5d**). Overall, we extend G9a function(s) beyond transcription to translational regulation during COVID-19 pathogenesis.

Emerging and future coronaviruses require new therapeutics to increase antiviral breadth, combat emerging drug resistance, and improve tolerability. Drugs that target G9a and its interacting protein EZH2 have known safety profiles and targeting methyltransferase complexes in a transient manner offers several advantages in the regulation of sarbecovirus infection. First, the translation regulatory function of G9a, by way of the G9a-METTL3-m^6^A axis, is anticipated to be synergistic with existing direct-acting antiviral (remdesivir, molnupiravir, paxlovid) and host immunomodulatory drugs (dexamethasone). Second, by suppressing secretion of ACE2 (**Figures 2d**), G9a inhibitors can hinder entry of ACE2-dependent viruses including SARS-CoV-1/2, HCoV-NL63, and various bat sarbecoviruses with potential for spillover to humans ^94–96^. Third, G9a targets multiple stages of the SARS-CoV-2 replication/life cycle, exhibiting multifaceted effects that include reduction in infection-induced hyperinflammation and thrombosis/lung-fibrosis marker expression, curbing T cell depletion/dysfunction and suppression of viral replication (**Figures 1d & 5d**; also see ref. ^97^). Fourth, the m^6^a modification pathway is hijacked by several RNA viruses (e.g., influenza A, HCV, HBV, EV71, HIV) for propagation and persistence ^98–101^ and, accordingly, sites of m^6^A modification are conserved in various SARS-CoV-2 variants ^80^, highlighting a potential evolutionary function of these sites in transmission and epidemiology. Indeed, Batista-Roche et al. reported that SARS-CoV-2 variants (epsilon > B.1.1.519 > alpha/gamma > omicron) and vaccination status (unvaccinated > partially-vaccinated > vaccinated) modulated genome and/or viral m^6^A levels differentially ^102^. Taken together, our data suggest that methyltransferase inhibitors (G9a/Ezh2) can be repurposed into broad-spectrum antivirals and represent a novel class of host mechanism-directed therapeutics to counter emerging drug resistance and infection. Comprehensive studies are needed in animal and/or human models (1) to further understand mechanistic details, (2) to evaluate efficacy and synergy of G9a/Ezh2 inhibition in combination with existing direct-action and immunomodulatory antivirals, and (3) to study potential antiviral activity against other m^6^A modification dependent RNA viruses.

Going forward, also needed is comprehensive investigation that discriminates between the canonical (transcriptional repression) and noncanonical (translation promotion) functions of the G9a complex during viral infection. Similarly, the exact function of EZH2 during the process requires further investigation. COVID-19 accelerates Alzheimer’s-related symptoms and dementia ^103, 104^, and we recently reported on development of a blood-brain-barrier penetrant G9a inhibitor that reversed cognitive and noncognitive effects of Alzheimer’s in multiple mouse models^45^. Thus, it will be interesting to study the effects of G9a inhibition in Alzheimer’s mouse models of COVID-19.

## ACKNOWLEDGMENTS

This work was supported in part by grants NIH R21AG071229, 1R41DK133051-01A1, R01 GM133107-01, and UNC University Cancer Research Fund (to X.C.). P.-Y. S. was partially supported by awards from the Sealy & Smith Foundation, the Kleberg Foundation, the John S. Dunn Foundation, the Amon G. Carter Foundation, the Summerfield Robert Foundation, and the Edith and Robert Zinn Foundation. NIH R35NS116843 (to H.S.), R01HD088626, R01AG084184, and an endowed professorship from the Icahn School of Medicine at Mount Sinai (to J.J.). NIH R01 NS125778 and John S. Dunn Research Foundation (to P.W.). NIH R35NS097370 and Dr. Miriam and Sheldon G. Adelson Medical Research Foundation (to G-l.M.). MS1262 is protected by US provisional patent applications jointly filed by Mount Sinai and UNC-CH. The indication of using clinically trialed G9a/Ezh2 inhibitors for COVID19 therapy is protected by US provisional patent application #63/113,211 as ‘Use of Method’.

## AUTHOR CONTRIBUTIONS

X.C. conceived and designed the project and experiments, secured funding, and wrote the manuscript.

A.M. designed and performed experiments, analyzed and interpreted data, and wrote the manuscript.

L. X. performed inhibitor treatment, sample preparation and processing for MS/MS experimental analysis and analyzed data.

F. Z., H. S., J.S. performed RNA-seq, MeRIP-seq, and analyzed the data.

X. X., P. S., P.W. performed inhibitor treatment and collected antiviral and cytotoxicity data.

J. A. W. analyzed RNA-seq, MeRIP-seq data.

Y. X., X.Y. and J.J. provided UNC0965, UNC0642, UNC1999 and YX59-126 targeting G9a/Ezh2.

## DECLERATION OF INTERESTS

The authors declare no competing interests.

## CONFLICT OF INTEREST STATEMENT

X.C. is the founder of TransChromix, LLC. J.J. is a cofounder and equity shareholder in Cullgen, Inc., a scientific cofounder and scientific advisory board member of Onsero Therapeutics, Inc., and a consultant for Cullgen, Inc., EpiCypher, Inc., Accent Therapeutics, Inc, and Tavotek Biotherapeutics, Inc. The Jin laboratory received research funds from Celgene Corporation, Levo Therapeutics, Inc., Cullgen, Inc. and Cullinan Oncology, Inc.

## MATERIALS AND METHODS

### Contact for Reagent and Resource Sharing

Further information and requests for resources and reagents should be directed to, and will be fulfilled by, the Lead Contact, Xian Chen.

## EXPERIMENTAL MODEL AND SUBJECT DETAILS

### Cell lines, patient samples, and treatment

Human alveolar epithelial cells that overexpress hACE2 receptor (A549-hACE2) [60] were maintained in high-glucose DMEM supplemented with 10% fetal bovine serum (FBS; HyClone Laboratories, South Logan, UT), 1% penicillin/streptomycin (P/S; 10,000 U/mL), 1% 4-(2-hydroxyethyl)-1-piperazineethanesulfonic acid (HEPES; ThermoFisher Scientific) and 10 μg/mL Blasticidin S (ThermoFisher Scientific). Peripheral blood mononuclear cells (PBMCs) isolated from patients with COVID-19 were purchased from RayBiotech Life (Peachtree Corners, GA) and were maintained in RPMI 1640 containing 10% FBS, 100 units/ml P/S, 2 mM L-glutaMax (Gibco), 10 mM HEPES and 1 mM Sodium Pyruvate at a density of 1.5×10^6^ cells/ml. The cells were treated with UNC0642, YX59-126, UNC1999 or TAZVERIK^®^ (Taz, Tazemetostat) at 1 µM for 0 to 48 h before being harvested for subsequent proteomic, RNA-seq, and molecular/cell biological studies. Cells were grown at 37 °C in humidified air with 5% carbon dioxide. All cell lines tested negative for mycoplasma.

## METHOD DETAILS

### Chemicals and reagents

Cell culture media, other components, and fetal bovine serum were obtained from Gibco. Trypsin was purchased from Promega. A549-hACE2 cell line was purchased from InvivoGen. All chemicals used during preparation of proteomic samples were HPLC-grade unless specifically indicated. Inhibitors (UNC0642, UNC1999 and MS1262) and probes (UNC0965, UNC2399) targeting G9a/Ezh2 were synthesized in Dr. Jian Jin’s lab. Tazemetostat (Cat. S7128) was purchased from Selleck. TMT 11-plex isobaric labeling reagent kit was purchased from Thermo-Fisher (Cat. A34808).

### SARS-CoV-2-Nluc antiviral assay

A549-hACE2 and SARS-CoV-2-Nluc were used for evaluating the compounds as described^105^. A549-hACE2 cells (12,000 cells per well in phenol red-free medium containing 2% FBS) were plated into a white opaque 96-well plate (Corning). On the next day, 3-fold serial dilutions of compounds were prepared in DMSO. One microliter of compound was mixed with 99 μl of SARS-CoV-2-Nluc virus that was diluted in phenol red-free culture medium containing 2% FBS. Compound-virus mixtures (50 μL) were added to each well of the 96-well plates containing A549-hACE2 cells (MOI 0.025). At 48 h post-infection, 50 μL of Nanoluciferase substrates (Promega) were added to each well. Luciferase signals were measured using a Synergy™ Neo2 microplate reader. The relative luciferase signals were calculated by normalizing the luciferase signals of the compound-treated groups to the signals of the DMSO-treated groups (set as 100%). The relative luciferase signal (Y-axis) versus the log10 values of compound concentration (X-axis) was plotted in software Prism 9. The EC_50_ (compound concentration for reducing 50% of luciferase signal) was calculated using a nonlinear regression model (four parameters). Two experiments were performed with technical duplicates.

### Cytotoxicity assay

A549-hACE2 cells (5000 cells per well in phenol red-free medium containing 2% FBS) were plated into a clear flat-bottom 96-well plate (Nunc). On the next day, 3-fold serial dilutions of compounds were prepared in DMSO. The compounds were further diluted 100-fold. Diluted compound solutions (50 μL) were added to each well of the cell plates. At 72 h post-treatment, 4 μL of Cell Counting Kit-8 (CCK-8; Sigma-Aldrich) was added to each well. After incubation at 37 °C for 90 min, absorbance at 450 nm was measured using the Cytation5 multi-mode microplate reader (BioTek). The relative cell viability was calculated by normalizing the absorbance of the compound-treated groups to the absorbance of the DMSO-treated groups (set as 100%). The relative cell viability (Y-axis) versus the log10 values of compound concentration (X-axis) were plotted in software Prism 8. The CC50 (compound concentration for reducing 50% of cell viability) was calculated using a nonlinear regression model (four parameters). Two experiments were performed with technical duplicates.

### m^6^A RNA immunoprecipitation sequencing (m6A/MeRIP-Seq) and data analysis

The m^6^A/MeRIP-Seq was performed as described with some modifications ^106^. Briefly, total RNA was extracted from A549-hACE2 and patient PBMCs using TRIzol (Life Technologies) followed by purification using illustra ^TM^ RNA spin Mini kit (GE Healthcare, UK). mRNA was isolated from 10 μg total RNA using Dynabeads Oligo (dT)_25_ (Thermo Fisher; 61006) according to manufacturer’s instructions. Ten percent of a total of 150 ng mRNA was used as input, while the rest was incubated with 3 ug anti-m^6^A polyclonal antibody (Synaptic Systems; 202003) that was pre-conjugated to Dynabeads Protein A (Thermo Fisher; 10001D) in 500 uL IP buffer (50 mM Tris, pH 7.4, 150 mM NaCl, 0.1% Igepal CA-630) for 2 hours at 4°C. After washing the beads twice with IP-buffer and twice with high-salt wash buffer (50 mM Tris pH 7.4, 500 mM NaCl, 0.1% Igepal CA-630) for 5 minutes each, the m^6^A-tagged mRNA was eluted using 100 uL IP-buffer containing 6.7 mM N^6^-Methyladenosine (Sigma-Aldrich; M2780) and 40 U RNase Inhibitor (NEB, M0314S) and then recovered with RNA Clean and Concentrator-5 spin columns (Zymo; R1015).

The input and m^6^A-IPed mRNA were subjected to library generation using the SMART-seq protocol as described^107^. For first strand cDNA synthesis, the mRNA was mixed with 0.25 µL RNase inhibitor and 1 µL CDS primer (5’-AAGCAGTGGTATCAACGCAGAGTACT30VN-3’) and heated to 70 °C for 2 min. Then the mixture containing 0.5 µL of 100 mM DTT, 0.3 µL of 200 mM MgCl_2_, 1 µL of 10 mM dNTPs, 0.25 µL RNase inhibitor, 1 µL of 10 µM TSO primer (5’-AAGCAGTGGTATCAACGCAGAGTACATrGrGrG-3’), 2 µL of 5X SMARTScribe RT buffer and 0.5 µL SMARTScribe reverse transcriptase (Takara, 639536) was added to perform reverse transcription. The cDNA was then amplified by Advantage Polymerase Mix (TAKARA, 639201) with IS primer (5’-AAGCAGTGGTATCAACGCAGAGT-3’). After purification with 0.8X AMPure XP beads (Fisher Scientific, A63880), the fragmentation of 100 pg cDNA was performed with EZ Tn5 Transposase (Lucigen, TNP92110). Fragments of cDNA were amplified by KAPA HiFi hotstart readymix (EMSCO/FISHER, KK2601) with the Nextera i7 primer and Nextera i5 primer. The DNA was purified with 0.8X AMPure XP beads and quantified by qPCR with KAPA Library Quantification Kit (Fisher, NC0078468). The DNA from different samples was pooled at equal molar amounts, and the final sequencing library was loaded at concentrations of 2.7 pM and sequenced on a NextSeq 550 (Illumina) for single-end sequencing.

Raw sequencing data were de-multiplexed using bcl2fastq2 v2.17.1.14 (Illumina) and adapters were removed using Trimmomatic-0.32 software. Then the Input and m6A-IP reads were mapped to a combined reference genome consisting of human (GRCh38/hg38) and Ensembl SARS-CoV-2 isolate Wuhan-Hu-1 genomes (Genome Assembly: ASM985889v3, Accession: GCA_009858895.3; Sequence: MN908947.3) using STAR v.2.7.6a. Only uniquely mapping reads at the exon level for each gene were quantified and summarized to gene counts. Differential gene expression analysis was performed using DESeq2 v.1.32.0. Resulting bam files were sorted and indexed using SAMtools v.1.1, and MACS2 v.2.2.7.1 was used for m^6^A peak calling using the bam files for each m^6^A-IP/input pair. The R packages GenomicRanges v1.36.1 and AnnotationHub v.3.4.0 were used to identify genes overlapping the peaks determined by MACS2, and CHIPseeker v.1.30.3 was used to create profile plots.

### ChaC pull-down with biotin-conjugated inhibitors UNC0965 and UNC2399

ChaC pull-downs were conducted as described^79^, with a few modifications. Briefly, cell pellets were resuspended in extraction buffer (50 mM Tris-HCl at pH7.5, 150 mM NaCl, 0.5% IGPAL-CA630 and 1 mM PMSF) followed by brief sonication and centrifugation to collect supernatant. Cell lysate (1 mg) was incubated overnight at 4°C with 2 nmole UNC0965/UNC2399 pre-coupled to 50 μl neutravidin-agarose (Thermo-Fisher) and washed thrice with 1 ml extraction buffer to remove nonspecific proteins. For on-beads sampling and processing, five additional washes with wash buffer (50 mM Tris-HCl pH7.5, 150 mM NaCl, 1 mM PMSF) were used to remove residual detergents. On-beads tryptic digestion was performed with 100 μl buffer containing 2 M urea, 25 mM Tris-HCl pH8.0, 1 mM DTT, 400 ng trypsin (Promega) for 30 min at room temperature on a mixer (Eppendorf). Partial digests were collected, and beads eluted twice more with 100 μl elution buffer (2 M urea, 25 mM Tris-HCl pH8.0, 5 mM iodoacetamide), incubated for 30 min at room temperature with shaking. Combined eluates were acidified with trifluoroacetic acid at final concentration of 1% (TFA, mass spec grade, Thermo-Fisher) and desalted with homemade C18 stage tips.

### Proteomics sample preparation

Patient PBMCs and A549-hACE2 cells were lysed in 2x Laemmli buffer followed by protein precipitation using a methanol-chloroform method ^108^. The resulting supernatant was mixed with Trizol LS reagent (Invitrogen, Cat 10296010), and proteins were further isolated and precipitated following the manufacturer’s instructions. Protein pellets were resuspended in 8 M urea, 50 mM Tris-HCl pH 8.0, reduced with dithiothreitol (5 mM final) for 30 min at room temperature, and alkylated with iodoacetamide (15 mM final) for 45 min in the dark at room temperature. Samples were diluted 4-fold with 25 mM Tris-HCl pH 8.0, 1 mM CaCl_2_ and digested with trypsin at 1:100 (w/w, trypsin:protein) ratio overnight at ambient temperature. Peptides were desalted using homemade C18 stage tips, and their concentrations were determined (Peptide assay, Thermo 23275). The cleaned peptides were used for LC-MS analysis or for additional labeling.

For TMT-labeling, 100 µg of each peptide sample was used for labeling with isobaric stable tandem mass tags (TMT11, Thermo Fisher Scientific, San Jose, CA) following manufacture instruction. The mixture of labeled peptides was desalted on Cep-Pak light C18 cartridge (Waters). Phosphopeptides were enriched with High-Select Fe-NTA Phosphopeptide Enrichment Kit (Thermo Scientific). Non-phosphopeptides (100 µg) were fractionated into 24 fractions using C18 stage tips and 10 mM TMAB (pH 8.5) containing 5-50% acetonitrile.

For secretome studies, secreted proteins were isolated from mixtures of culture supernatant and Trizol LS reagent (Invitrogen, Cat 10296010) following manufacturer’s instruction. Resulting protein pellets were solubilized in buffer containing 50 mM Tris (pH 8.0) and 8 M urea, then processed in the same way as that for proteome to obtain purified peptides. LFQ was used for protein quantification of secretome.

### LC-MS/MS analysis details parameters

Dried peptides were dissolved in 30 μl of 0.1% formic acid, 2% acetonitrile. One microgram phosphopeptides or 0.5 μg non-phosphopeptides from each fraction was analyzed on a Q-Exactive HF-X coupled with an Easy nanoLC 1200 (Thermo Fisher Scientific, San Jose, CA). Peptides were loaded on to a nanoEase MZ HSS T3 Column (100 Å, 1.8 µm, 75 µm x 250 mm, Waters). Parameters for MS and LC are listed in **supplemental table S6**.

### Raw proteomics data processing and analysis

Mass spectra were processed, and peptide identification/quantification performed using MaxQuant software version 2.1.0.0 (Max Planck Institute, Germany). All protein database searches were performed against the UniProt human protein sequence database (UP000005640). False discovery rates (FDR) for peptide-spectrum match (PSM) and protein assignment were set at 1%. Search parameters included up to two missed cleavages at Lys/Arg, oxidation of methionine, protein N-terminal acetylation and phosphorylation on serine, threonine, and tyrosine as dynamic modifications. Carbamidomethylation of cysteine residues was considered as a static modification. Peptide identifications are reported by filtering of reverse and contaminant entries and assigning to their leading razor protein. Data processing and statistical analysis were performed using Perseus (Version 1.6.10.50). Protein quantitation was performed on biological replicates and a two-sample t-test statistic was used with a p-value of 5% to report statistically significant protein or phosphopeptide abundance fold-changes.

### Analysis of functional category and networks

Canonical pathway, biological function and upstream regulator analyses were performed using IPA (https://www.ingenuity.com/), DAVID (http://david.abcc.ncifcrf.gov/), Metascape (https://metascape.org/), and STRING (http://string-db.org/). Figures were generated using RStudio and interactome analyses were performed in Cytoscape v3.8.0.

## SUPPLEMENTAL MATERIALS

**Supplemental Figure S1:**
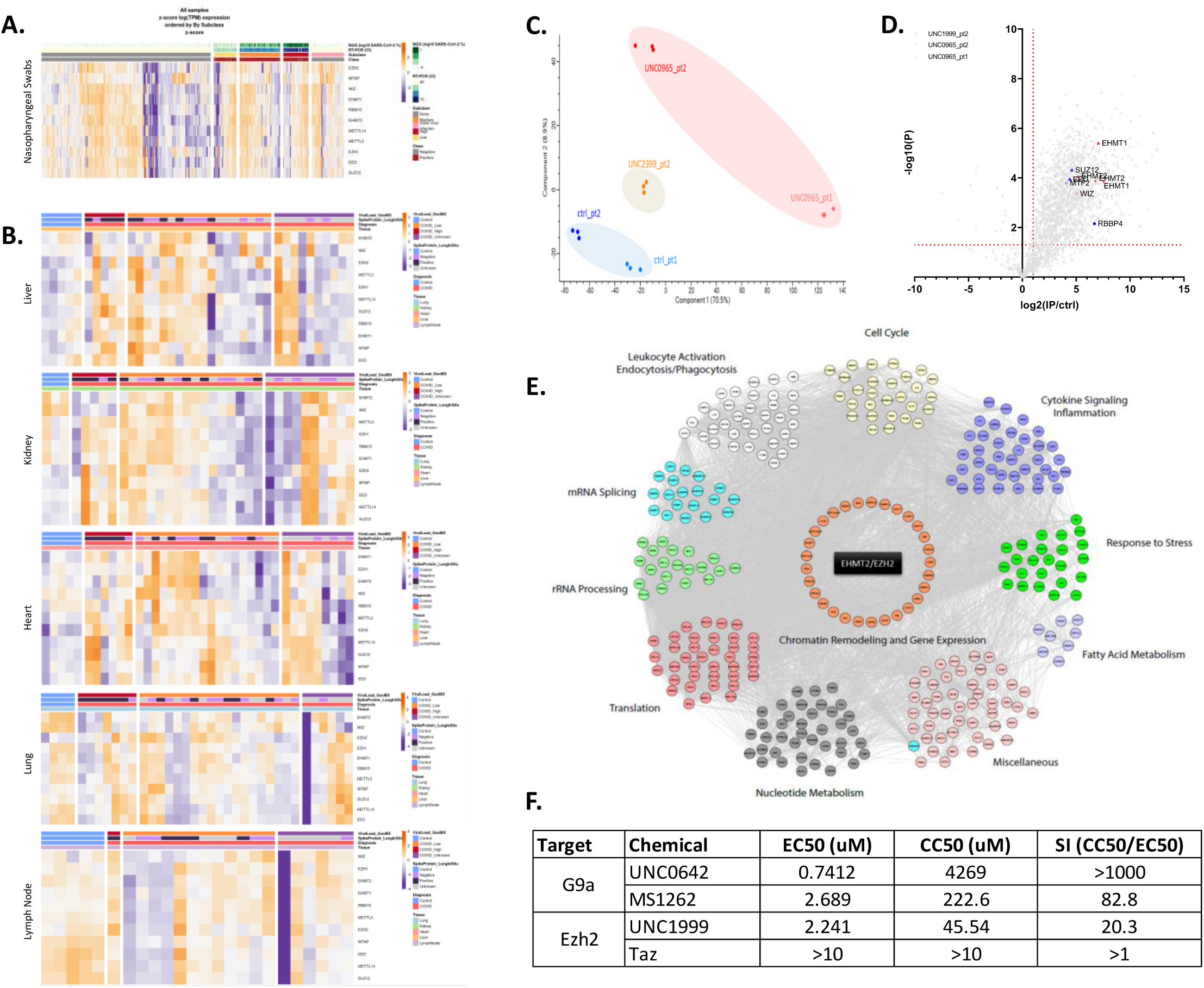
G9a, Ezh2, and METTL3 complex members are upregulated in COVID-19 patient samples and other ChaC-MS/antiviral-assay related figures. (**A**) Heatmap depicting overexpression of G9a, EZH2, and METTL3 complex members in nasopharyngeal swabs from COVID-19 patients with high viral load. Patient samples have been stratified into ‘none’, ‘low’, ‘medium’, and ‘high’ based on viral load as defined by presence of SARS-CoV-2 RNA (detected using NGS and qRT-PCR from nasal swabs). ‘Other viral infection’ category includes patients with unrelated viral infections and serves as an additional control. Members that undergo significant (FDR corrected P < 0.01) transcriptional regulation in response to SARS-CoV-2 infection are shown. (** Figure generated using data from ref. ^109^). **(B)** Heatmaps depicting overexpression (RNA-Seq) of G9a, EZH2, and METTL3 complex members in autopsy samples (liver, kidney, heart, lung, lymph node) from COVID-19 patients. Patients were stratified into ‘control’, ‘COVID_Low’, ‘COVID_High’, and ‘COVID_unknown’ based on spatial transcriptomics (GeoMx) and expression (in situ hybridization against viral S gene) profiling of autopsy samples. Members that undergo significant (FDR corrected P < 0.01) transcriptional regulation in response to SARS-CoV-2 infection are shown. (** Figures generated using data from ref. ^110^). **(C)** Principal Component Analysis (PCA) plot shows clear separation between UNC0965 (G9a, 2 patients), UNC2399 (Ezh2, 1 patient), and control (2 patients) pulldowns from COVID-19 patient-derived PBMCs. Experiments were performed in triplicate. **(D)** Volcano plot showing enrichment of G9a/Ezh2 interactors (log2(FC) > 1; p-value < 0.05) from COVID-19 patient-PBMCs with core G9a/Ezh2 complex components highlighted. **(E)** Protein-protein interaction map of G9a/Ezh2 shared interactors (368) identified using ChaC-MS from COVID-19 patient-derived PBMCs. In addition to chromatin-remodeling/transcription related proteins, G9a/Ezh2 complexes interact with translation regulatory, cell cycle, immune response, and RNA metabolism related proteins, among others, during Covid19 pathogenesis. **(F)** Summary of antiviral/cytotoxicity assay results for compounds targeting G9a and Ezh2 in human A549-hACE2 cells. (See also Figure 1**, Supplemental Table S1**)

**Supplemental Figure S2:**
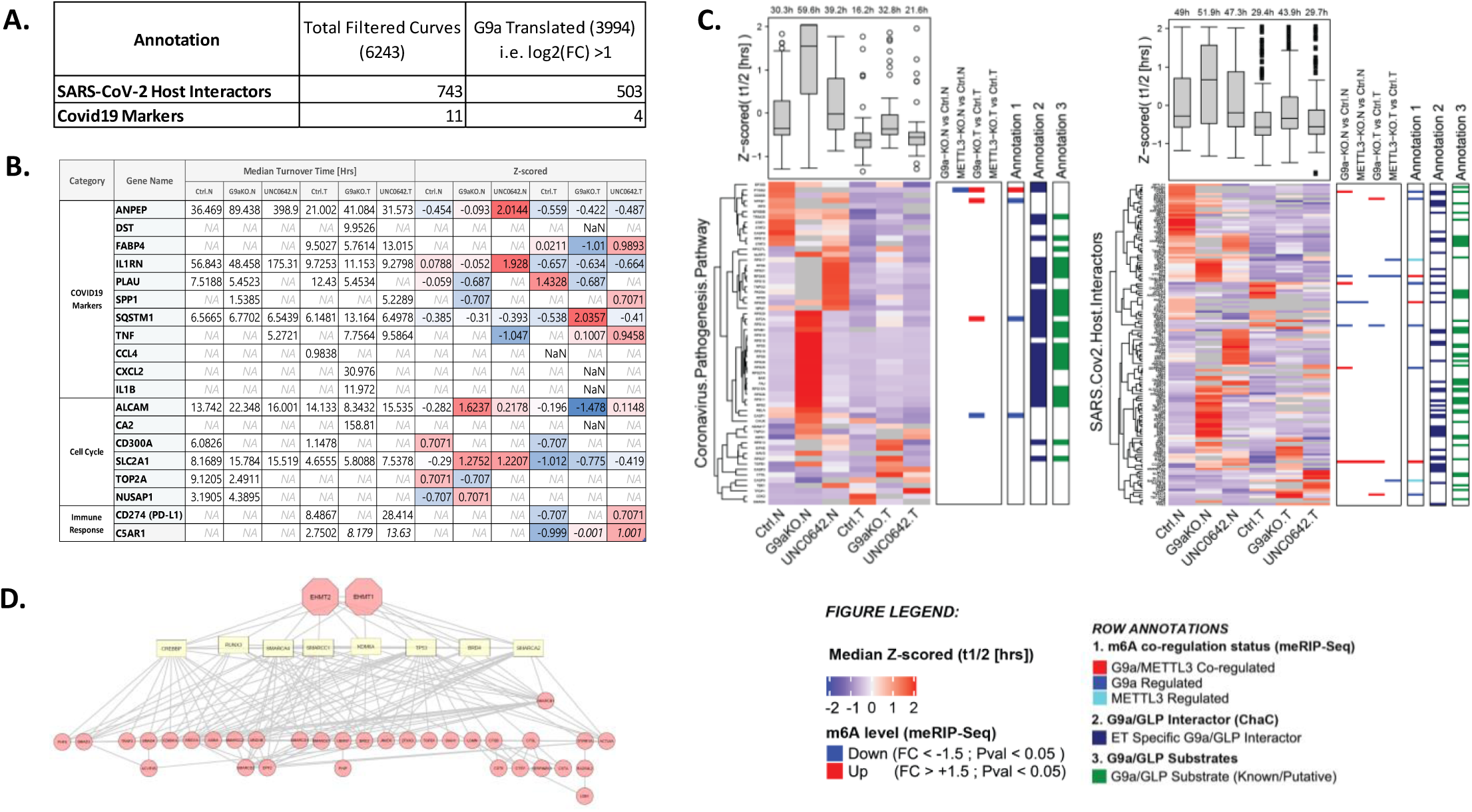
G9a promotes turnover of SARS-CoV-2 pathogenesis-related proteins and loss/inhibition of G9a and its associated proteins hinders SARS-CoV-2 replication and infection. (**A**) G9a loss/inhibition affects turnover of numerous host interactors of SARS-CoV-2-encoded proteins and COVID-19 markers in murine (Raw 264.7) ET macrophage cells. **(B)** Table showing effect of G9a loss/inhibition on select proteins of interest. **(C)** Heatmaps showing that ET promotes turnover of numerous host interactors of SARS-CoV-1/2– and MERS-encoded proteins and coronavirus pathogenesis pathway related factors, whereas G9a loss/inhibition reverses effect of ET on turnover of these coronavirus related proteins. **(D)** Genetic perturbation of several G9a-associated proteins adversely affects SARS-CoV-2 replication and infection. **Note:** Panels (A-C) generated using data from ref. ^79^ whereas (D) adapted from refs. ^21, 62^). (See also **Supplemental Table S1**).

**Supplemental Figure S3:**
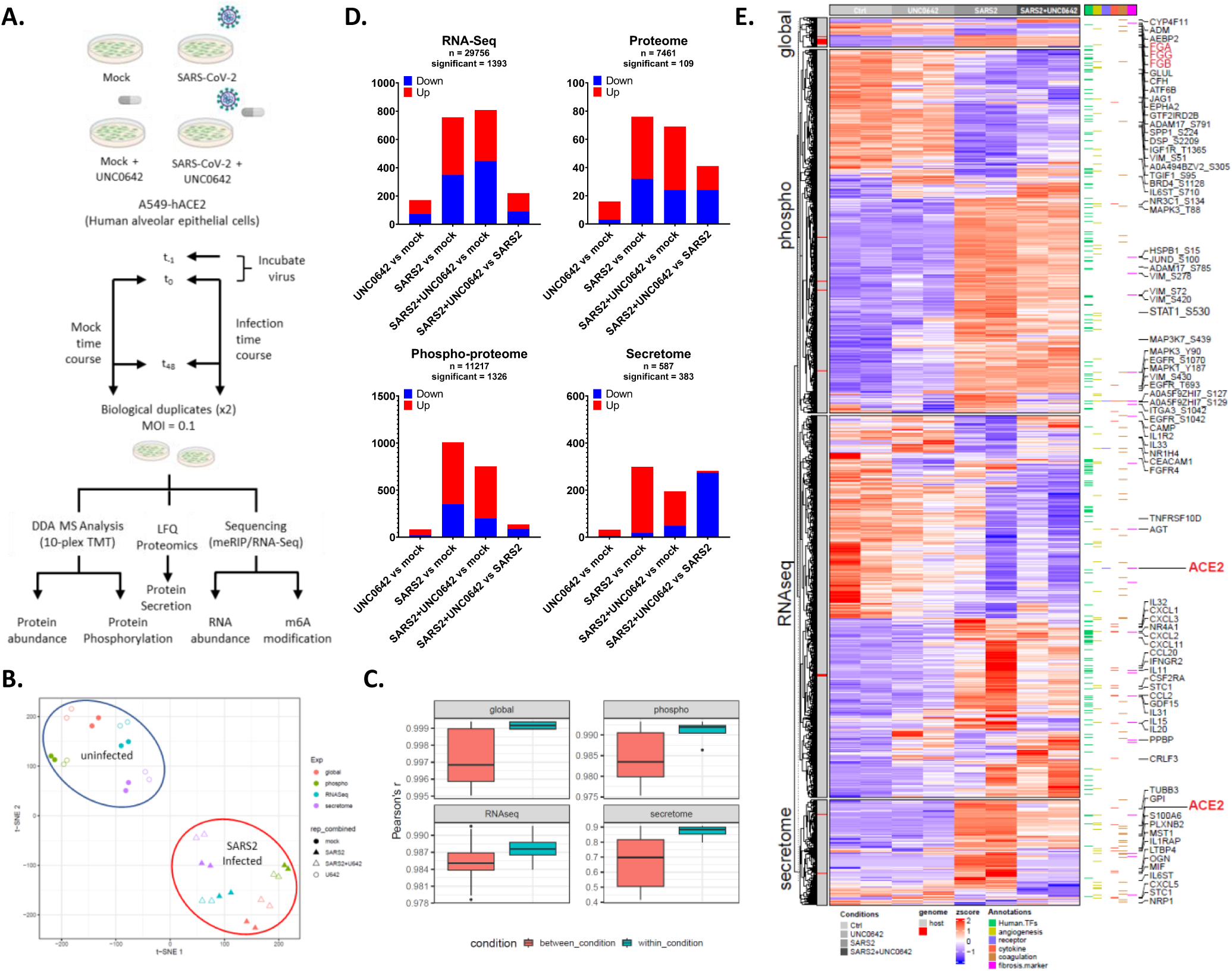
Multi-omic dissection of G9a’s function during SARS-CoV-2 infection. (**A**) A549-hACE2 cells were infected with SARS-CoV-2 (MOI = 0.1). After 1 h of virus uptake, media were switched, and cells/media were harvested (48 hpi) with/without UNC0642 treatment. As a control, A549-hACE2 cells were also mock infected for 1 h and harvested after 48 h of mock infection. All experiments were performed in biological duplicate. Following cell/media harvest, RNA was collected for m^6^A/RNA-Seq, and proteins were digested into peptides followed by mass spectrometry (MS) to measure changes in protein abundance upon infection; the remaining sample was enriched for phosphorylated peptides and analyzed to measure changes in phosphorylation signaling. A 10-plex TMT based DDA approach was used for all MS acquisitions to detect global and phospho-proteomic changes, although an LFQ approach (with 3 technical replicates each) was used to detect SARS-CoV-2 mediated changes to secretome. **(B)** The t-distributed stochastic neighbor embedding (t-SNE) visualization of significantly dysregulated proteins in tested conditions (shape) and datasets (color). **(C)** Correlation between replicates within a biological condition (red) and across biological conditions (green). Boxplots depict median (horizonal lines), interquartile range (boxes), maximum and minimum values (vertical lines), and outliers (solid circles). **(D)** Bar plots depict number of dysregulated entities (red = up; blue = down) for indicated comparisons in RNA-Seq, proteome, phospho-proteome, and secretome datasets from A549-hACE2 cells. Total number of identified (n) and dysregulated (significant) entities is mentioned on top. **(E)** Heatmap of dysregulated transcripts (1393 transcripts), proteins (109 proteins), phospho-sites (1326 PHs), and secreted proteins (383 proteins) in SARS2/mock infected A549-hACE2 cells with/without UNC0642 treatment. Entities are clustered according to dataset, viral and host entities are indicated (left annotation column) and six functional clusters of SARS-CoV-2 pathogenesis relevant viral receptors and proteases, transcription factors, cytokines/chemokines, coagulation system-related proteins, angiogenesis-associated proteins, and fibrosis markers are shown (right annotations). Names of select receptors and cytokines/chemokines are also highlighted. **(F)** Gene Ontology enrichment analysis for differentially regulated entities identified in (D-E). Top 20 terms with significant over-representation (adjusted P < 0.05) are shown, and redundant terms are removed. Grey cells indicate the lack of significant enrichment. **(G)** Circos plots show overlap among differentially regulated entities identified in (D-E). Purple lines represent shared genes between various datasets (left), whereas overlaid blue lines represent the different genes that fall in the same ontology term (right). **(H)** Protein-protein interaction enrichment analysis results for SARS2-dysregulated/UNC0642-reversed host entities shown in Figure 2D. Briefly, molecular complex detection (MCODE) algorithm was used to identify densely connected network components followed by application of pathway/process enrichment analysis to each MCODE component independently. Best representative term(s) were retained as functional description of the corresponding components. Primary receptor used by SARS-CoV-2 (ACE2) is highlighted. **(I)** Heatmap showing that SARS-CoV-2 infection affects expression/secretion/phosphorylation of various m^6^A readers/writers/erasers, an effect that was largely reversed following UNC0642 treatment of infected A549-hACE2 cells. (See also Figures 2 **& 3, Supplemental Table S2**)

**Supplemental Figure S4:**
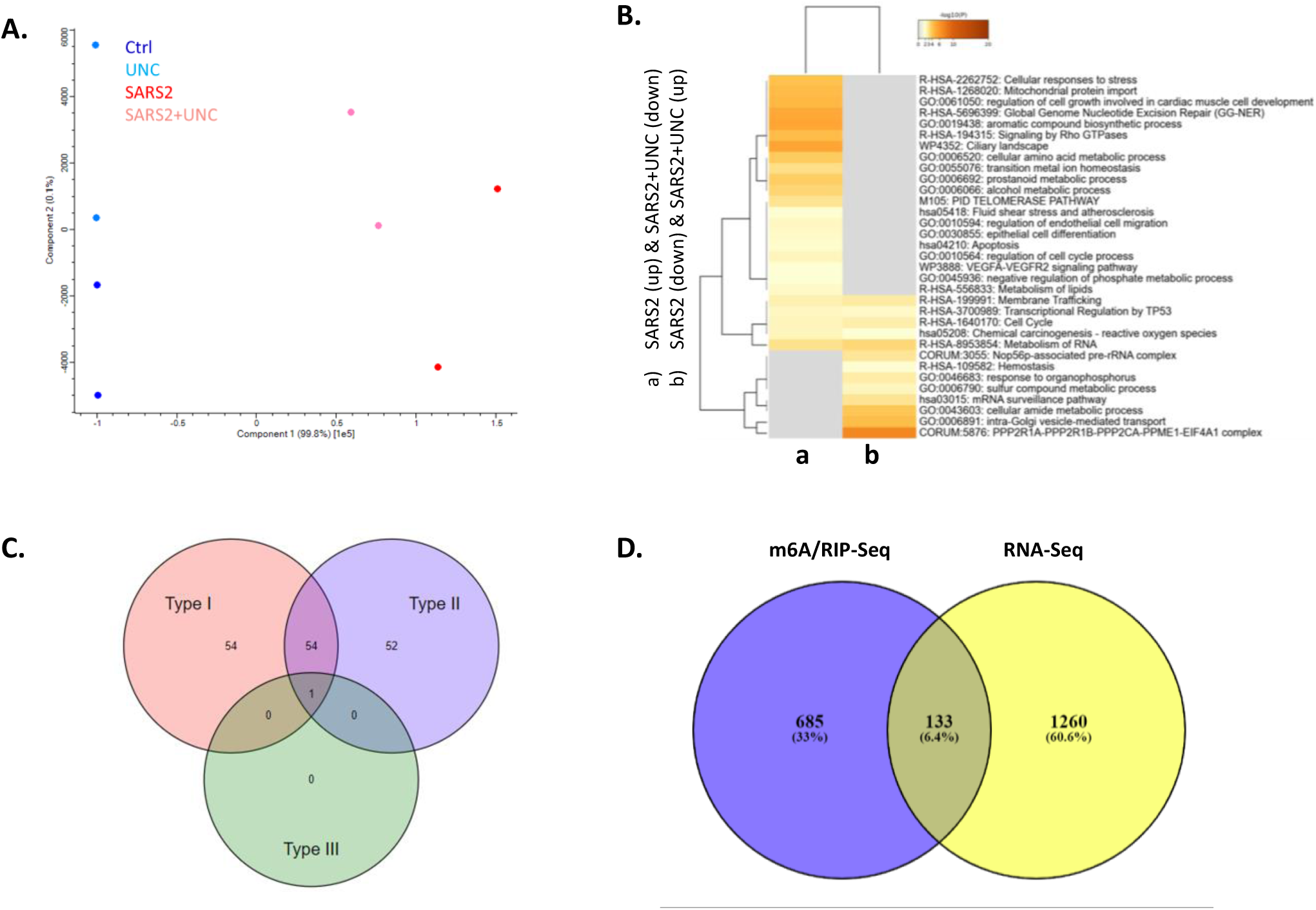
G9a promotes m6A modification of immune response related transcripts following SARS-CoV-2 infection without altering mRNA expression. (**A**) PCA plot showing that SARS-CoV-2 infected A549-hACE2 cells have distinct m6A epi-transcriptomic landscape compared to uninfected controls. Similarly, UNC0642/DMSO treated samples cluster separately from each other. **(B)** Pathway enrichment analysis for SARS-CoV-2 dysregulated, UNC0642 reversed, m6A modified transcripts identified using meRIP-Seq in A549-hACE2 cells. Top20 terms with significant over-representation (adjusted P < 0.05) are shown, and redundant terms are removed. Grey cells indicate the lack of significant enrichment. **(C)** Venn diagram showing number/type of interferon regulated genes (IRGs) among SARS-CoV-2 dysregulated, UNC0642 reversed, m6A transcripts identified using meRIP-Seq in A549-hACE2 cells. Plot generated using interferome (http://www.interferome.org/) **(D)** Venn diagram showing that most transcripts with differential m6A modification (meRIP) did not exhibit corresponding change in mRNA expression (RNA-Seq). (See also Figure 4**, Supplemental Table S3**)

**Supplemental Figure S5:**
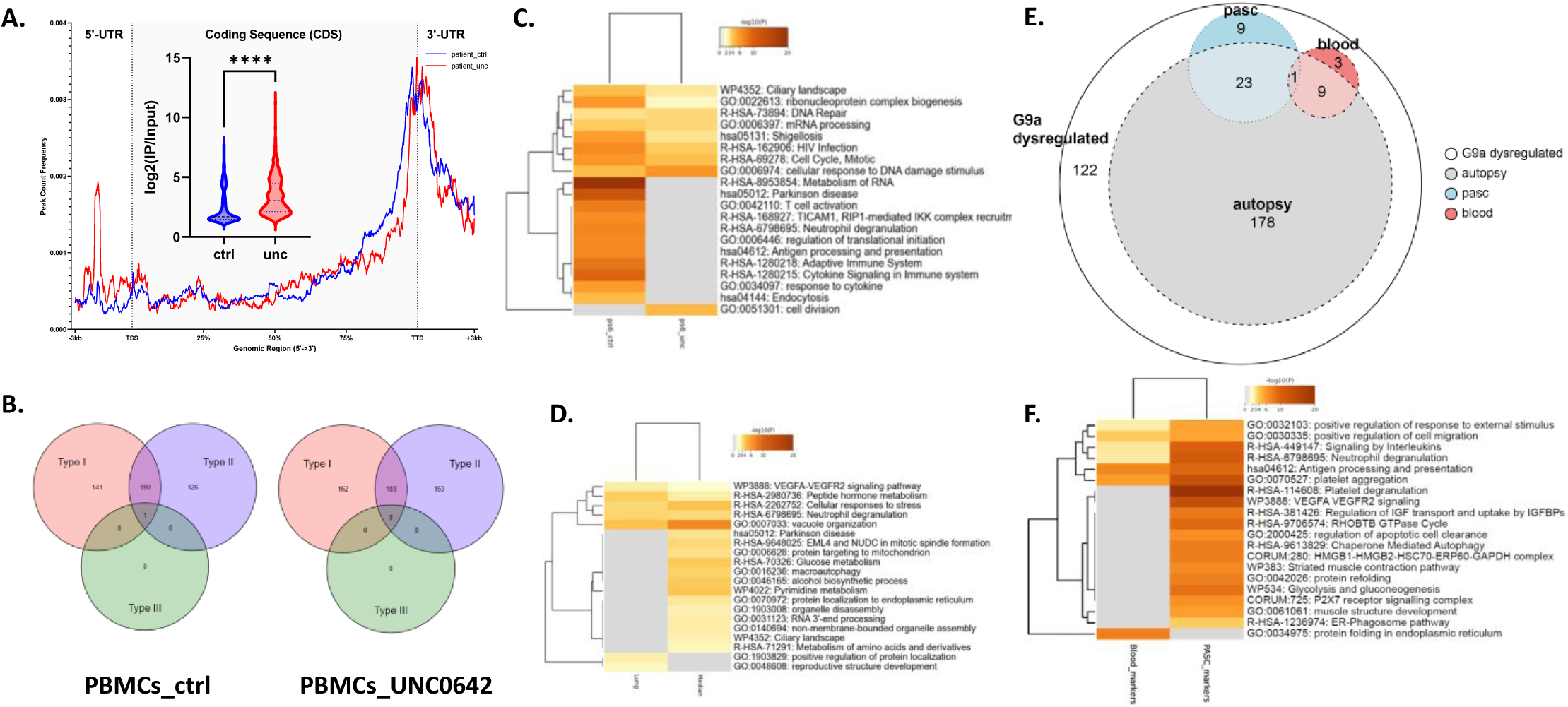
UNC0642 inhibition affects m6A modification of immune response related transcripts in patient PBMCs. (**A**) Profile plot showing distribution of the enriched m6A peaks in control (blue line) and UNC0642 treated (red line) COVID-19 patient-derived PBMCs analyzed along the RNA segments. Each transcript was length normalized and ±3Kb from TSS/TTS are included. Violin plot shows an increase in m6A level following UNC0642 treatment (red), compared to DMSO treated controls. Mann-Whitney test was used for statistical analysis (****p < 0.0001). **(B)** Venn diagrams showing number/type of interferon regulated genes (IRGs) among m6A modified transcripts identified in patient PBMCs. Plots were generated using interferome (http://www.interferome.org/). **(D)** Pathway enrichment analysis for SARS-CoV-2 dysregulated, UNC0642 reversed, poised-mRNAs identified in A549-hACE2 cells that showed similar dysregulated pattern in COVID-19 patient autopsy samples (shown in main Figure 5c). Top20 terms with significant over-representation (adjusted P < 0.05) are shown, and redundant terms are removed. Grey cells indicate the lack of significant enrichment. **(E)** Venn diagram showing overlap between SARS2-dysregulated/G9a-reversed proteins shown in Figure 2d and markers of long-covid (PASC) ^81^, blood markers of severe COVID19 ^82, 83^, and proteins dysregulated in autopsy samples from COVID19 patients^111^. **(F)** Pathway enrichment analysis for SARS2-dysregulated/G9a-reversed markers of severe (blood) and long (PASC) covid shown in (E). Top20 terms with significant over-representation (adjusted P < 0.05) are shown, and redundant terms are removed. Grey cells indicate the lack of significant enrichment. (See also Figure 6**, Supplemental Table S5**)

**Supplemental Figure S6:**
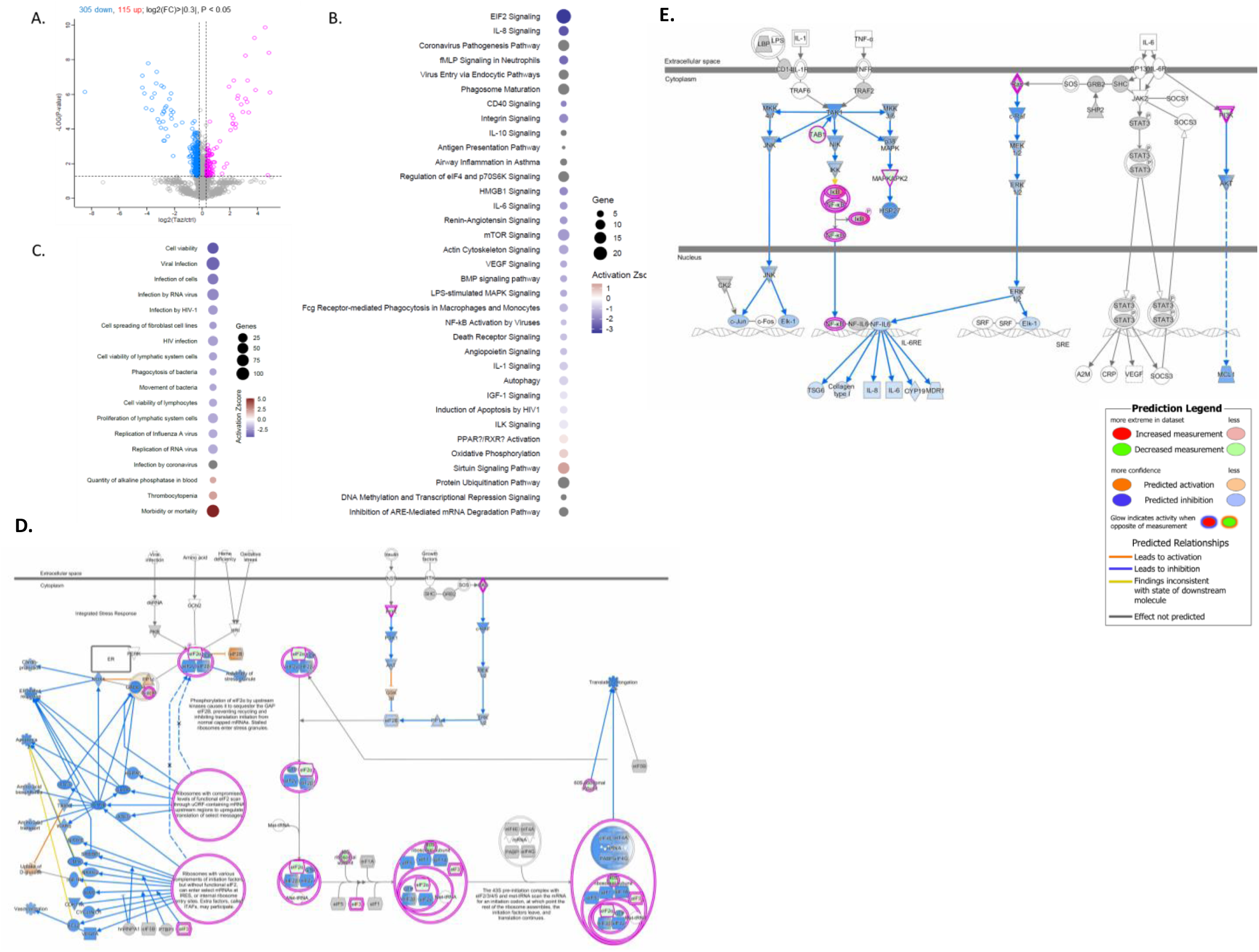
Ezh2 inhibition in Covid19 patient derived PMBCs leads to downregulation of viral infection related signaling and pathways. (**A**) Tazemetostat treated (1uM for 24h) PBMCs from severe Covid19 patients were pooled, fractionated, and subjected to LC-MS analysis. Each sample was run in triplicate. Volcano plot shows statistically significantly proteins (log2(FC)>|0.3|; Student’s t-test: P < 0.05) showing increased (red) and decreased (blue) expression respectively when compared to mock (DMSO) treated PBMC cells. **(B-C)** Significantly altered signaling cascades **(B)** and disease related pathways **(C)** upon tazemetostat treatment are shown (Fisher’s Exact test; adjusted P < 0.05). Size of the dot represents number of significantly altered genes belonging to the pathway while color represents Z-score representing activation (red) or deactivation (blue) of said pathway. Pathways for which activation/deactivation could not be inferred are shown in grey. (**D-E**) These networks overlaid with protein expression changes following tazemetostat treatment of patient PBMCs show reduction in activity of translation (EIF2/4) and inflammation (NF-kB, IL-6, IL-8) related pathways, compared to DMSO treated controls (i.e., UNC1999 vs DMSO). **Note:** Ingenuity Pathways analysis (IPA) was used to generate these networks. Nodes colored red/green represent increased/decreased expression in our dataset, while orange/blue colored nodes & edges represent predicted activation/deactivation of said regulators based on our data. Yellow color means that observation in our dataset is inconsistent with literature while grey color means that activity could not be predicted. (See also **Supplemental Table S5c**)

**Supplemental Table S1:** Summary of patient-PMBC/ET-macrophage ChaC-MS results along with translatome & meRIP-Seq data for coronavirus pathogenesis related proteins from ET macrophage cells.

**Supplemental Table S2:** Summary of multiomic analysis of SARS-CoV-2/mock infected A549-hACE2 cells with or without UNC0642 treatment.

**Supplemental Table S3:** Summary of multiomic data to identify differentially m6A modified transcripts (meRIP-Seq) and poised-mRNAs (RNA-Seq, LFQ-MS) in SARS-CoV-2/mock infected A549-hACE2 cells with or without UNC0642 treatment.

**Supplemental Table S4:** Summary of multiomic data (proteomics, RNA/meRIP-Seq) to identify poised-mRNAs in COVID-19 patient-derived PBMCs & correlated analysis of G9a regulated poised-mRNAs in multi-organ autopsy samples.

**Supplemental Table S5:** Summary of global/phospho-proteomic data from UNC1999 treated mock/SARS2 infected A549-hACE2 cells. Analysis of tazemetostat treated COVID-19 patient PBMCs is also included.

**Supplemental Table S6:** Details of LC-MS parameters for different global/phospho/secretome datasets.

## Notes

### Competing Interest Statement

The authors have declared no competing interest.

